# Statistical analysis of repertoire data demonstrates the influence of microhomology in V(D)J recombination

**DOI:** 10.1101/2024.10.16.618753

**Authors:** Magdalena L Russell, Assya Trofimov, Philip Bradley, Frederick A Matsen

**Author notes:** To whom correspondence should be addressed. (M.L.R.) or (F.A.M.). **Author contributions:** M.L.R., A.T., P.B., and F.A.M. designed research; M.L.R. performed research; M.L.R. and F.A.M. analyzed data and wrote the paper.

## Abstract

V(D)J recombination generates the diverse B and T cell receptors essential for recognizing a wide array of antigens. This diversity arises from the combinatorial assembly of V(D)J genes and the junctional deletion and insertion of nucleotides. While previous *in vitro* studies have shown that microhomology—–short stretches of sequence homology between gene ends—–can bias the recombination process, the extent of microhomology’s impact *in vivo*, particularly in humans, remains unknown. In this paper, we assess how germline-encoded microhomology influences trimming and ligation during V(D)J recombination using statistical inference on previously-published high-throughput TCR*α* repertoire sequencing data. We find that microhomology increases both trimming and ligation probabilities, making it an important predictor of recombination outcomes. These effects are consistent across different receptor loci and sequence types. Further, we demonstrate that accounting for microhomology effects significantly alters sequence annotation probabilities and rankings, highlighting its practical importance for accurately inferring the V(D)J recombination events that generated an observed sequence. Together, these results enhance our understanding of how microhomologous nucleotides shape the human V(D)J recombination process.

**Significance Statement:** Humans rely on diverse adaptive immune receptor repertoires to effectively defend against infections. The receptor generation process, known as V(D)J recombination, is designed to create this diversity by stochastically joining V(D)J gene segments and modifying their junctions through nucleotide deletions and insertions. Previous studies, conducted in vitro, have suggested that short stretches of homologous nucleotides between gene segments can bias these recombination steps. In this study, we explore the extent to which these homologous nucleotides influence V(D)J recombination in humans using statistical inference on large-scale receptor repertoire sequencing data. Our findings reveal that microhomology significantly biases several recombination steps, with important practical implications for the analysis, processing, and interpretation of receptor sequences.

## Introduction

V(D)J recombination is an essential process for generating diverse B cell receptors (BCRs) and T cell receptors (TCRs). In this process, single V-, D- (if present), and J-genes are randomly selected from a pool of germline gene segments, then edited and joined together to form a uniquely recombined receptor sequence. Previous *in vitro* experiments have suggested that short stretches of sequence homology between gene ends, known as microhomology, can play a significant role in the V(D)J recombination process. This raises the question of whether microhomology impacts V(D)J recombination *in vivo*, particularly in terms of recombination outcomes in humans with intact recombination machinery. Understanding this has practical implications for V(D)J recombination sequence *annotation*. Annotation means inferring the specific V(D)J recombination editing and joining processes that produced each sequence, forming the basis for many downstream B cell and T cell repertoire analyses. In this paper, we use statistical inference on high-throughput human TCR repertoire data to assess how microhomology influences various steps of the V(D)J recombination process.

In order to more fully set the stage, we will now summarize the relevant biological context. V(D)J recombination begins when the recombination activating gene (RAG) protein complex aligns two randomly chosen genes, removes the intervening chromosomal DNA between the two genes, and forms a hairpin loop at the end of each gene [15, 12, 43]. Each hairpin loop is then nicked open by the Artemis:DNA-PKcs complex [31, 29, 43]. Hairpin opening most frequently occurs at position +2, where position 0 refers to the edge of the hairpin and position −1 refers to the last nucleotide on the 5’ strand [31], however, other hairpin opening positions are also possible [31, 29]. The Ku heterodimer (Ku70/Ku80) can bind to each nicked gene end and recruit non-homologous end joining factors, in any order, to repair the double stranded break [30, 27, 28]. From here, it is likely that the processing of the two gene ends occurs iteratively, with multiple rounds of action by a nuclease, polymerase, and ligase which eventually leads to a joining event to combine the two gene fragments [28, 38].

The various possible processing steps involved in this iterative end-joining stage are as follows. Nucleotides can be deleted from each gene end through a mechanism suggested to involve the Artemis nuclease [11, 34, 35, 24, 19, 37, 9, 6, 46, 42]. Nucleotide deletion is thought to occur in a sequence-dependent fashion; for example, sequences with high AT content have been found to experience greater nucleotide loss than those with high GC content [34, 14, 35, 24], and the extent of deletions has been shown to depend on local nucleotide identity [33, 41], as well as sequence breathing capacity and length [41]. Additionally, non-template-encoded nucleotides, known as N-insertions, can be added by terminal deoxynucleotidyl transferase (TdT) [25, 17, 26]. TdT has a bias for the addition of purine-purine and pyrimidine-pyrimidine di-nucleotides suggesting that nucleotide addition depends on the previous addition [14, 33]. Further, nucleotide addition lengths and composition have been shown to depend on the presence (or absence) of nucleotide trimming at the gene ends [13]. Joining of the two gene ends is then carried out by XRCC4:DNA ligase IV, a flexible ligase that can ligate across gaps and incompatibilities between the ends, along with additional end-joining factors like XLF and PAXX that stabilize the ends, and polymerases that fill in gaps [20, 21, 1, 36, 8].

The presence of microhomology, while not required, has been suggested to bias the outcome of the random V(D)J recombination processing steps. Microhomology can occur in several forms: (1) **terminal microhomology**^1^, found at the ends of genes and encoded in the germline; (2) **interior microhomology**, located *within* the sequences and also germline-encoded; and (3) **insertion-dependent microhomology**, created by N-insertions and not encoded in the germline. If present, terminal microhomology can directly guide ligation without additional processing. In contrast, interior and insertion-dependent microhomology may necessitate deletions or further N-insertions before microhomology-mediated ligation can occur.

Experiments *in vitro* and with model organisms have suggested that microhomology (i.e. 1-4 nucleotides) is an important factor in V(D)J recombination. Although microhomology between gene ends is not essential for joining (Figure 1A part (i)) [16, 4], it has been shown to improve joining efficiency and bias the outcome towards using the microhomologous region to guide trimming and ligation (Figure 1A part (ii)) [14, 30, 28, 37, 8, 7, 38]. For example, reconstitution experiments suggest that sequences with microhomology can stabilize gene ends without requiring additional end-joining factors like XLF and PAXX and germline-encoded microhomology may reduce the necessity for template-independent addition by polymerase-*µ* and TdT [8], possibly explaining the enhanced ligation efficiency. *In vitro* studies show that 1 or 2 nucleotides of germline-encoded microhomology are present in nearly 60% of ligated coding joints in the absence of TdT [14], with similar observations reported in neonatal mice when TdT levels are low [10, 18]. However, this frequency drops substantially when TdT is present, as TdT-mediated additions are thought to create stronger insertion-dependent microhomology [14, 28, 37]. The involvement of microhomology in ligation appears to be more complex when it is not present at sequence ends or generated through nucleotide addition. Most gene ends lack terminal microhomology after hairpin opening but share interior microhomology [29, 8, 7]. In such cases, the Artemis–DNA-PKcs complex has been shown to trim gene ends to expose interior microhomology (Figure 1A part(ii)) [8, 7].

**Figure 1:**
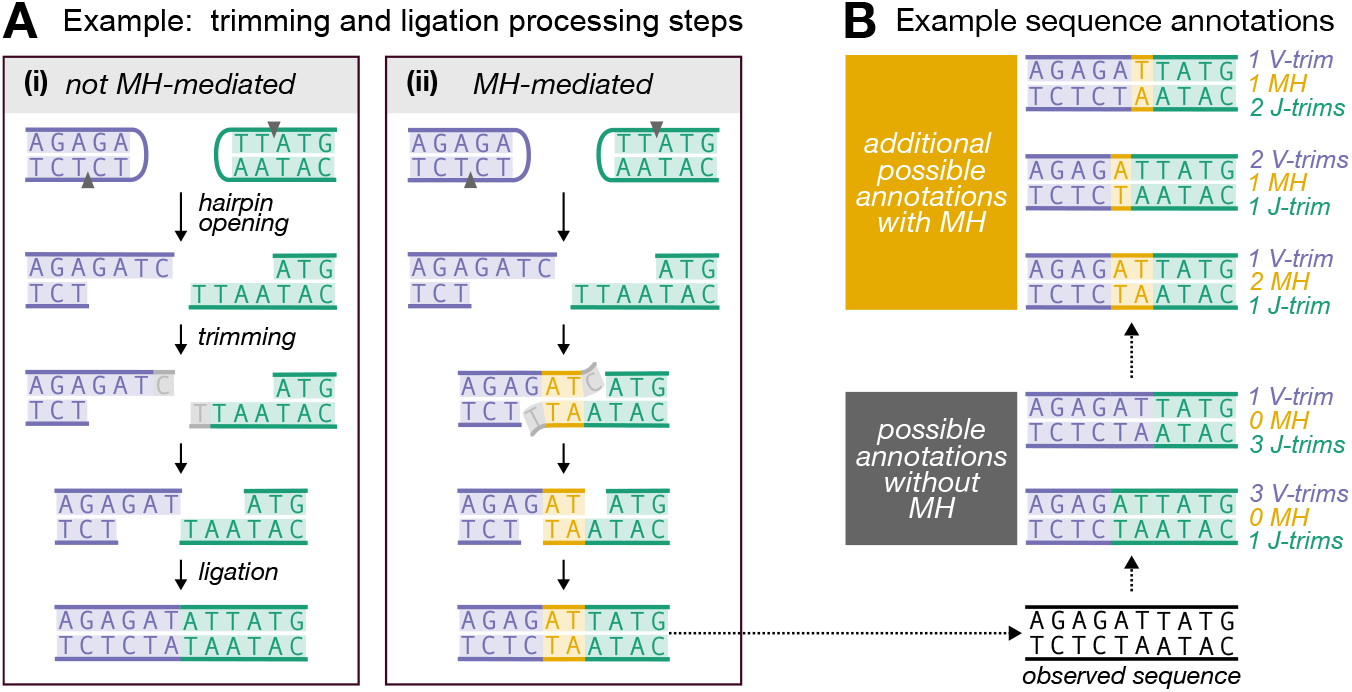
**(A)** Illustration of how germline microhomology (MH) could affect trimming and/or ligation during V(D)J recombination. We use sequences without N-insertions to quantify these effects, leveraging germline V- and J-gene sequences to identify potential MH-mediated ligation events. The example shows microhomologous regions (yellow) and trimmed nucleotides (gray) for a V-gene (purple) and J-gene (green), highlighting two regimes: (i) no MH influence and (ii) MH-mediated trimming and ligation. MH could affect trimming, ligation, or both, leading to distinct sequences regardless of whether the genes are trimmed equally or differently. **(B)** Illustration of possible V(D)J recombination annotations for sequences potentially ligated with MH and lacking N-insertions. Existing annotation software does not account for MH and assigns shared nucleotides to only one sequence (gray box annotations). However, additional MH annotations (yellow box) are possible but not considered. Our modeling aims to include all annotations, both with and without MH, for each sequence.

This essential biochemical work has demonstrated that microhomology can significantly affect V(D)J recombination, however, it does not demonstrate its importance for shaping V(D)J recombination in humans. In addition to being an issue of intrinsic interest, the role of microhomology has practical implications as well: if microhomology impacts the probability of V(D)J recombination annotations (i.e. numerical histories of recombination events such as gene choice, trimming, insertion, ligation, etc.), then corresponding terms should be incorporated into software that infers recombination probabilities. This would ensure that additional annotations involving microhomology are also considered (Figure 1B).

Statistical inference on high-throughput repertoire sequencing datasets allows exploration of the *in vivo* V(D)J recombination mechanism in humans. In fact, existing probabilistic models of V(D)J recombination, such as IGoR [32], have provided interesting and important insights about the natural underlying mechanism by learning statistics of V(D)J recombination. These models have revealed significant dependencies between recombination events, such as gene usage and trimming, and have provided estimates of the overall probabilities of generating specific TCR sequences, thereby helping to disambiguate the effects of generation from selection [33, 32]. Similar statistical approaches have been successfully applied to understand the sequence-dependent process of nucleotide trimming, revealing significant connections between trimming patterns and local sequence identity, length, and wider GC content [41]. However, to our knowledge, no probabilistic models of V(D)J recombination incorporating microhomology have been developed.

In this paper, we explore the extent to which germline-encoded microhomology biases trimming and ligation during V(D)J recombination using statistical inference on high-throughput TCR*α* repertoire sequencing data [23, 22]. We have designed a flexible probabilistic modeling framework, allowing us to quantify the extent to which microhomology biases trimming and ligation probabilities. Our results show that the presence of microhomology significantly increases trimming and ligation probabilities, and is an important predictor of the choices made in these processes. These observations are consistent with sequences from an independent TCR*α* validation dataset, as well as with sequences from other receptor loci such as TCR*γ*. Additionally, we demonstrate that explicitly including microhomology-related terms in our model significantly impacts sequence annotation probabilities and overall V(D)J recombination annotation rankings. Together, these findings enhance our understanding of microhomology’s involvement in the V(D)J recombination process and highlight the importance of accounting for microhomology-related effects in receptor sequence processing and analysis.

## Results

### Data and data processing overview

We analyzed TCR*α*-immunosequencing data from 10 individuals [23, 22]. The *TRA* locus was chosen for its higher sequence diversity between joining genes (V- and J-gene pairs) compared to the *TRB* locus. Each sequence was assigned with a V(D)J recombination annotation using the IGoR software, which is designed to learn unbiased recombination statistics from immune sequence reads [32]. IGoR generated a list of potential recombination annotations with corresponding likelihoods for each sequence in the training dataset. We then assigned each sequence with a single annotation by sampling from these possible annotations according to their posterior probabilities.

We excluded sequences with N-insertions to leverage our knowledge of germline V- and J-gene sequences in identifying potential germline-microhomology-mediated ligation events. N-insertions complicate the analysis of ligation patterns because the composition of inserted nucleotides before ligation is unknown, and their presence suggests that *germline*-microhomology-mediated ligation did not occur.

Since IGoR does not consider microhomology and assigns shared nucleotides to only one sequence, we adapted the initial IGoR-inferred annotations to account for microhomology. This adaptation extended the range of possible V(D)J recombination annotations for each sequence, resulting in a set of possible microhomology-adapted annotations for each sequence (see Figure 1B and SI Appendix for details).

Additionally, TCR sequences can be categorized as “productive” if they code for a functional protein, or “non-productive” otherwise, arising from out-of-frame recombination or presence of stop codons. Each T cell can undergo recombination at two alleles; if the first is non-productive and the second successful, both sequences can be sequenced as part of the repertoire. Non-productive sequences do not generate proteins for thymic selection, and their recombination statistics should reflect only the V(D)J recombination process [40, 33, 44]. In contrast, productive sequence statistics reflect both recombination and selection. To study nucleotide trimming and ligation during V(D)J recombination without selection effects, we included only non-productive sequences in our training dataset.

### Terminology

In this paper, we discuss the mechanisms of trimming and ligation as they occur between V- and J-gene pairs during V(D)J recombination. Below, we define key terms used throughout the paper:

- **Trimming scenario**: A specific pair of trimming events, one at the V-gene end and one at the J-gene end.
- **Ligation scenario**: A specific number of microhomologous nucleotides shared between the trimmed V-gene and J-gene, facilitating their ligation. The possible ligation scenarios for a given V-J gene pair are determined by their germline sequences and the extent of trimming.
- **Joint trimming and ligation scenario probability**: The normalized probability of a particular combination of trimming and ligation scenarios occurring for a V-J gene pair, considering all possible combinations for that pair.
- **V(D)J recombination annotation**: A specific set of V(D)J recombination events that produce a sequence, including trimming, insertion, and ligation scenarios.
- **V(D)J recombination annotation probability**: The normalized probability of a particular V(D)J recombination annotation for an observed sequence, calculated from all possible annotations for that sequence.

Since our analysis is restricted to sequences without N-insertions, we derive these probabilities from joint trimming and ligation scenario probabilities and normalize over all possible scenario combinations for that specific observed sequence.

### Methods overview

In our previous work, we established that local nucleotide identities at trimming sites (the “trimming motif”) and the counts of GC or AT nucleotides beyond these motifs (the “two-side base-count beyond”) are strong predictors of trimming probabilities for single gene sequences [41]. Building on this foundation, we have integrated these established parameters with newly developed microhomology-related parameters to assess their combined effects on trimming and ligation processes.

To this end, we developed a two-step conditional logit model to evaluate the joint probabilities of trimming and ligation scenarios for V- and J-gene pairs. The model describes a generative process in two steps:

1. **Trimming scenario choice**: This choice is determined by the established “trimming motif” and “two-side base-count beyond” parameters for each gene, in addition to a new parameter that quantifies the effect of microhomology on trimming. Specifically, this parameter measures the importance of the average number of microhomologous nucleotides between two trimmed sequences, a value that varies depending on the chosen trimming scenario.
2. **Ligation scenario choice**: This choice is determined by a novel microhomology parameter related to ligation, which quantifies the importance of the number of microhomologous nucleotides that ultimately appear in the final ligated sequence.

The mental model of this two-step process is that trimming occurs first, independently of ligation, and then ligation occurs, conditioned on the trimming scenario.

These parameters are designed to quantify how sequence-level features, particularly microhomology, influence the decisions made during V(D)J recombination. Importantly, the magnitude of microhomology’s influence in guiding these choices is quantified by these model parameters, highlighting its role in the recombination process. We validated the model’s ability to detect these effects through a series of simulations (see SI Appendix). In order to assess the significance of microhomology-related terms in downstream analyses, such as in V(D)J recombination sequence annotation, we designed the model with the flexibility to include or exclude microhomology-related parameters for both trimming and ligation decisions.

For model training, we used non-productive sequences without N-insertions, along with corresponding sets of potential microhomology-adapted trimming and ligation scenarios for each sequence (as detailed in the previous section). We assigned probabilities to each potential scenario based on their likelihood from our model. Because these probabilities depend on the model’s parameters, we employed an expectation-maximization algorithm for parameter inference, which converged within twenty-five iterations (Figure S1). Further details about this algorithm can be found in the Methods and SI Appendix sections.

### Microhomology significantly increases trimming and ligation probabilities

Complementary sequence regions capable of forming microhomologous regions during V(D)J recombination are common between germline V- and J-genes in the *TRA* locus. The median average number of microhomologous nucleotides across the ensemble of possible trimming scenarios for these germline V- and J-gene pairs is 0.1978 (Figure S2). This median corresponds to 1.3149 possible ligation scenarios per trimming scenario (Figure S3 and Figure S4). Given that a median of exactly one ligation scenario per trimming scenario would indicate all V(D)J recombination annotations involve zero microhomology, this suggests that many trimming scenarios allow for multiple ligation outcomes, both with and without microhomology. Additionally, complementary sequence regions and their corresponding ligation scenario options are distributed across trimming scenarios depending on the specific V- and J-gene pair (Figure S2 and Figure S4). This distribution highlights the potential for both interior and terminal microhomology to influence trimming and ligation outcomes.

To quantify the effects of microhomologous nucleotides on trimming and ligation, we employed our model, which incorporates various sequence-level parameters, including those related to microhomology. We validated the model’s capability to detect microhomology effects through a series of simulations, designed to generate sequences by sampling trimming and ligation scenarios under different microhomology regimes: no microhomology effect, microhomology affecting either trimming or ligation choices exclusively, and microhomology influencing both. After training our model using each of these simulated datasets, we confirmed its sensitivity to detecting variable microhomology effects across different conditions (Figure S5).

We then applied our model to the real TCR*α* training dataset to quantify the actual effects of microhomology, along with other sequence-level features, on trimming and ligation probabilities. Each model parameter reflects the change in the log_10_ odds of trimming and/or ligating at a specific scenario due to an increase in the corresponding feature value, assuming all other features remain constant. We assessed the significance of each model parameter’s influence on trimming and ligation probabilities by estimating their standard errors with bootstrap methods and applying a z-test to obtain a p-value (see Methods). We used a Bonferroni-corrected significance threshold of 0.0016, adjusted for the total number of model parameters, and report parameters on the log_10_ scale. Our results indicate that the number of microhomologous nucleotides between two sequences substantially influences both trimming (parameter = 0.4484) and ligation (parameter = 0.1272), with both effects being highly significant (p-values smaller than machine tolerance, *p* ≃ 0) (Figure 2A). This relationship is further demonstrated by notable increases in joint trimming and ligation probabilities for scenarios with more microhomology, as illustrated in Figure 2B, which highlights two trimming and ligation scenarios from the most common V-J gene pair, TRAV41*01 and TRAJ45*01. While the influence of microhomology on trimming was stronger than on ligation, these effects appear to be interdependent. Interestingly, when training the model using sequences containing N-insertions (indicating a lack of germline microhomology-dependent ligation), microhomology had a small but significant effect on trimming probabilities (parameter = 0.0059; p-value smaller than machine tolerance, *p* ≃ 0) (Figure S6). This demonstrates that microhomology can independently influence trimming, suggesting a nuanced role beyond its interaction with ligation.

**Figure 2:**
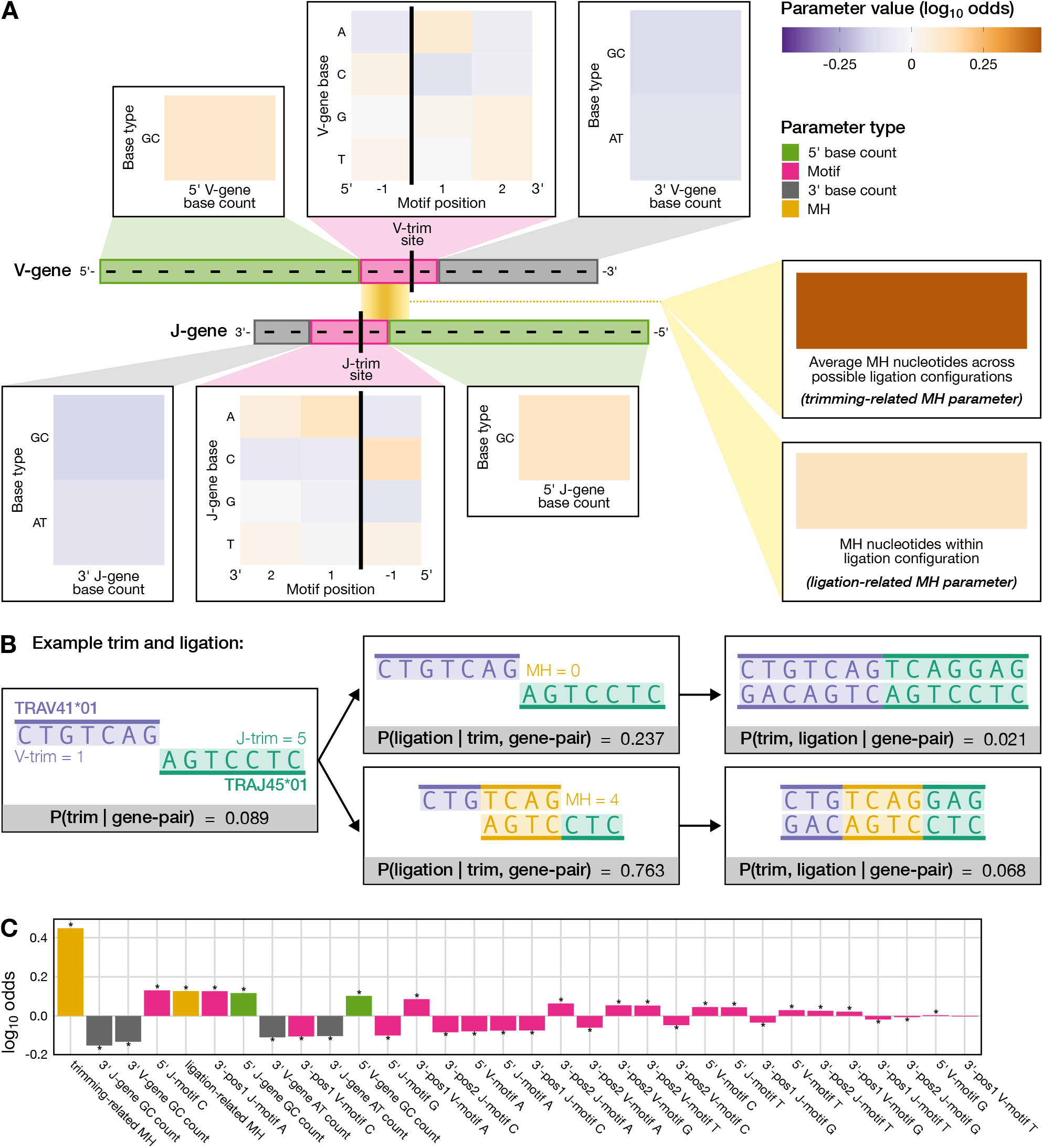
Although sequence-based parameters such as gene-specific trimming motifs and base counts contribute meaningfully to predicting trimming and ligation probabilities, the extent of microhomology (MH) between sequences exerts a strong effect, especially in increasing trimming probabilities. **(A)** Inferred microhomology (MH) parameters for trimming and ligation, along with V- and J-gene-specific trimming motif and base count parameters, alongside an illustration of how these sequence features align with an arbitrary V- and J-gene pair. V- and J-gene trimming motif parameters (pink) reflect the influence of adjacent nucleotides on trimming probabilities. For V-gene and J-gene 5’ and 3’ base counts (green and gray), an increase in GC nucleotides 5’ of the motif increases trimming probabilities, while an increase in AT or GC nucleotides 3’ of the motif decreases them. The model excludes 5’ AT nucleotide counts. MH between sequences (gold box) significantly influences both trimming and ligation probabilities, with a stronger effect on trimming. Black vertical lines indicate example trimming sites. Each parameter represents the change in log_10_ odds of trimming or ligating due to an increase in the feature value, assuming all other features are held constant. **(B)** Our model with inferred coefficients shows that increasing MH generally raises both trimming and ligation probabilities, as shown in example trimming and ligation scenarios for the most frequently used gene pair, TRAV41*01 (purple) and TRAJ45*01 (green). The bottom row shows the most probable trimming and ligation scenario (left and middle boxes), using four nucleotides of MH (gold), with a joint probability of 0.068 (right box). The top row shows that the same trimmed sequences could ligate with zero MH, leading to a lower joint probability of 0.021. **(C)** Both MH-related parameters (gold) exhibit large effect sizes, with the trimming-related MH parameter showing the strongest positive effect. Stars indicate parameters significant at a Bonferroni-corrected threshold of 0.0016.

Returning to the original model, in addition to microhomology effects, we identified significant “trimming motif” and “two-side base count” parameters for both V- and J-gene trimming probabilities. These parameters, previously introduced in our analyses of trimming patterns for single V- and J-gene sequences [41], showed results consistent with our previous work. As in our prior work, the local sequence context (“trimming motif”) for each gene was modeled using a position weight matrix from a three-nucleotide window around each trimming site. We observed similar patterns for both V-gene and J-gene local trimming contexts, where C and A nucleotides had the largest influence on trimming probabilities (Figure 2A). The “two-side base count” parameters include 5’ and 3’ base count parameters for each gene. The 5’ base count parameters act as a proxy for sequence-breathing effects, capturing preferences for GC content upstream of the motif, while the 3’ base count parameters reflect preferences for both the absolute position of the trimming site and sequence-breathing effects related to AT and GC content downstream of the motif. Our analysis showed that increasing GC nucleotides 5’ of the motif (which decreases sequence-breathing capacity) raised trimming probabilities. In contrast, increasing both AT and GC nucleotides 3’ of the motif (which increases absolute position) reduced trimming probabilities for both gene types (Figure 2A).

Finally, we examined the relative effect sizes of these sequence-level parameters to identify the most influential factors for trimming and ligation probabilities (Figure 2C). The strongest positive impacts were observed for trimming-related microhomology effects (parameter = 0.4484), followed by the presence of a C nucleotide immediately 5’ of the J-gene trimming site (parameter_*J*_ = 0.1308), and ligation-related microhomology effects (parameter = 0.1272). In contrast, the most negative influences were an increase in GC nucleotides 5’ of the motif for both V- and J-genes (parameter_*V*_ = −0.3043, parameter_*J*_ = −0.3482), the presence of a C nucleotide 3’ of the V-gene trimming site (parameter_*V*_ = −0.1049), and an increase in AT nucleotides 5’ of the motif for both V- and J-genes (parameter_*V*_ = −0.1095, parameter_*J*_ = −0.1030). P-values corresponding to each of these effects were smaller than machine tolerance (*p* ≃ 0).

### Microhomology is an important predictor of trimming and ligation probabilities across receptor loci

To assess the importance of incorporating microhomology-related parameters for accurately predicting trimming and ligation probabilities, we compared the performance of a full model, which includes microhomology, motif, and 5’ and 3’ base count terms, to models lacking specific terms. All models were trained using the non-productive TCR*α* training dataset and the parameters were held constant for subsequent analyses.

We began by evaluating model performance on the training dataset. The full model showed a substantially lower expected per-sequence log loss compared to the model without microhomology-related parameters, indicating a better fit to the data (Figure 3A). This improvement was validated by a likelihood ratio test (LRT), which confirmed the statistical significance of including microhomology terms (LRT statistic = 93754.84; p-value less than machine tolerance, *p* ≃ 0). The full model also exhibited higher predictive accuracy, as indicated by a lower mean absolute error (MAE) (MAE = 0.00468) compared to the model without microhomology terms (MAE = 0.00481). We repeated this analysis across models lacking other parameter types and found that the full model consistently outperformed them, exhibiting lower expected per-sequence log loss and MAE in each case (Figure 3). Recall that the 5’ base count parameters capture potential sequence-breathing effects by reflecting preferences for GC content upstream of the motif, while the 3’ base count parameters capture both preferences for the absolute position of the trimming site and sequence-breathing effects related to AT and GC content downstream of the motif. Among the individual parameter types, the 3’ base count terms had the largest impact, leading to the greatest improvement in both log loss and MAE. Microhomology and motif terms contributed the second-largest improvements in MAE and log loss, respectively. That is, the absolute position of the trimming site, represented by the 3’ base count terms, had the strongest influence, while the local nucleotide context at the trimming site (captured by motif terms) and the extent of microhomology between the trimmed and ligated sequences also provided positive contributions, though to a lesser extent. Sequence-breathing capacity upstream of the trimming site, reflected by the 5’ base count terms, improved log loss and MAE as well, but had a smaller overall effect compared to the other parameters.

**Figure 3:**
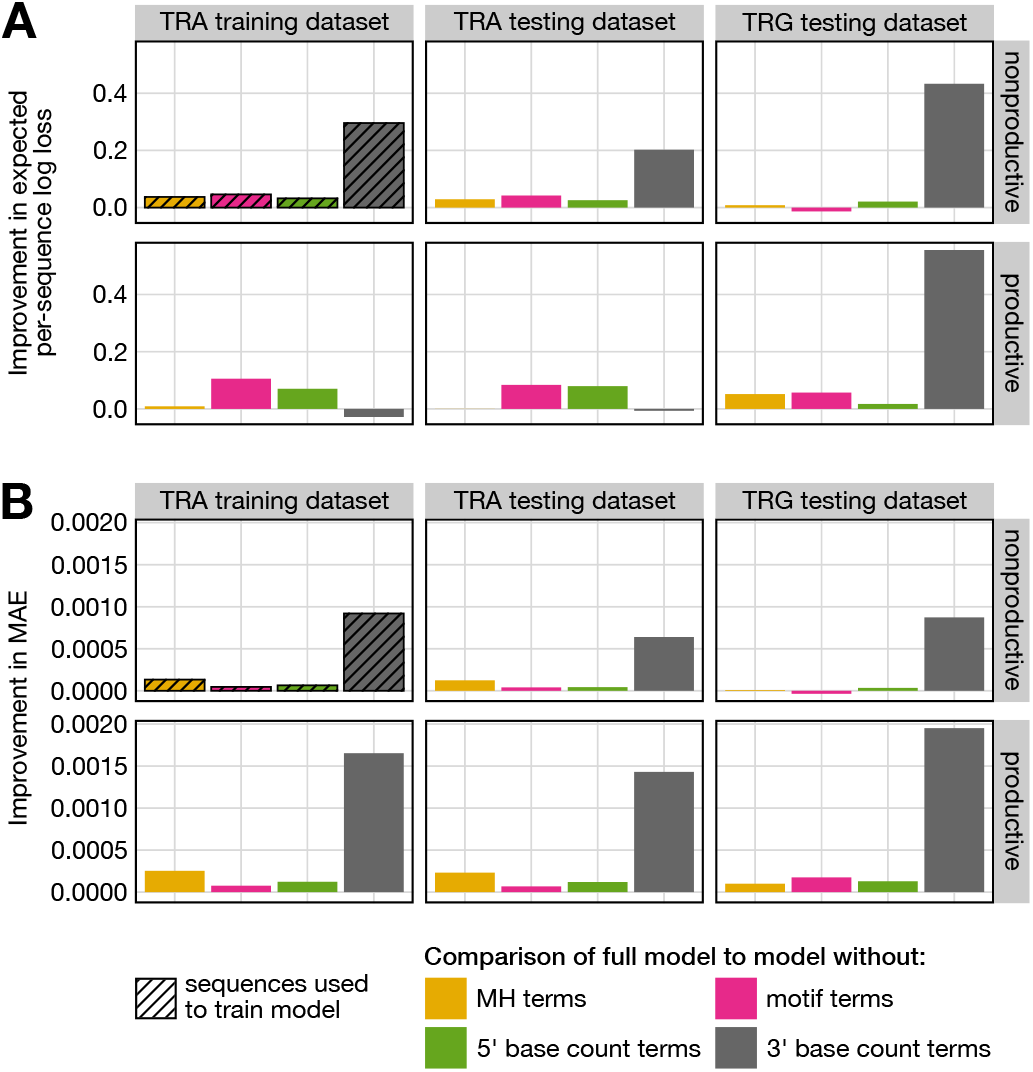
**(A)** Improvement in expected per-sequence log loss for the full model, which includes microhomology (MH) terms, motif terms, and 5’ and 3’ base count terms, compared to models without specific terms across both productive and nonproductive sequences from multiple datasets. Improvement is the negative difference in log loss, with negative values indicating a relatively worse fit and positive values indicating a relatively better fit for the full model. Including MH terms improves log loss across all datasets, except for productive sequences from the TCR*α* testing dataset, where no change in loss was observed. **(B)** Improvement in mean absolute error (MAE) across the same models and datasets. Improvement is the negative difference in MAE, with negative values indicating relatively lower predictive accuracy and positive values indicating relatively higher predictive accuracy for the full model. Including MH terms consistently improves MAE across all datasets. All models were trained using nonproductive sequences from the TCR*α* training dataset (hatched boxes), with parameters held constant (“frozen”) before calculating log loss and MAE across datasets.

Next, we assessed model performance using both productive and nonproductive sequences from various testing datasets, including independent TCR*α* and TCR*γ* datasets. The full model consistently demonstrated superior predictive accuracy across both productive and nonproductive sequences, with lower expected per-sequence log loss and MAE compared to the other models (Figure 3).

In most datasets, the inclusion of 3’ base count terms continued to have the strongest impact on improving model fit and predictive accuracy. However, there were two notable exceptions in log loss calculations for productive sequences from the TCR*α* training and testing datasets. In these cases, including 3’ base count terms, which capture effects related to the absolute positioning of the trimming site, negatively affected log loss. Since productive sequences are subject to selection-related effects that may alter preferences for trimming site positioning, the 3’ base count terms learned from nonproductive sequences—–where these selection effects are absent–—may be less effective for predicting trimming in productive sequences. Nevertheless, the inclusion of 3’ base count terms still improved MAE in these cases, despite the negative impact on log loss. This discrepancy may stem from log loss being more sensitive to outliers than MAE.

The inclusion of microhomology terms also improved model fit and predictive accuracy across most datasets, consistently providing the second-largest improvement in MAE. Notably, even when applied to productive sequences from the TCR*α* testing set–—despite these sequences not being included in training and having skewed recombination statistics due to selection—–the full model outperformed the model without microhomology terms in MAE, although the log loss values were similar. This suggests that while the inclusion of microhomology terms improves log loss across datasets, their most pronounced impact is on MAE. Overall, these consistent findings across different receptor loci and sequence types highlight the biological significance of microhomology in accurately modeling trimming and ligation scenarios.

### Accounting for microhomology affects sequence annotation

Given the significant role of microhomology in predicting trimming and ligation scenarios across receptor loci, we wanted to evaluate how microhomology parameterization influences sequence annotation. Recall that sequence annotation involves assigning a specific V(D)J recombination annotation, which describes the associated trimming, insertion, and ligation scenarios, to an observed sequence. In earlier sections, we examined the joint probabilities of trimming and ligation scenarios *for V-J gene pairs*, which represent the normalized probability of each trimming and ligation scenario within the complete set of possibilities for a given gene pair. Here, we shift our focus to V(D)J recombination annotation probabilities *for individual observed sequences*, which represent the normalized probability of each V(D)J recombination annotation within all possible annotations for a given sequence. Since we are analyzing sequences without N-insertions, each V(D)J recombination annotation corresponds directly to a trimming and ligation scenario, allowing us to use our inferred joint trimming and ligation scenario distributions to calculate the corresponding V(D)J recombination annotation probabilities. In this analysis, we compare the V(D)J recombination annotation probabilities and rankings between two models: (1) the full model, which includes microhomology, motif, and 5’ and 3’ base count terms, and (2) a version of the model that excludes microhomology terms.

As expected from our earlier results, we find that accounting for microhomology effects substantially changes annotation probabilities and their rankings. In fact, an average of 38.07% of gene pair sequences that have multiple possible V(D)J recombination annotations exhibit a different top-ranked annotation when using the model that parameterizes microhomology to calculate annotation probabilities compared to using the model that does not (Figure 4A). The extent of this affect appears to depend on the amount of potential microhomology present. Specifically, as the average microhomology across potential annotations increases for a given sequence within a V-J gene pair, so does the proportion of gene pair sequences with differing top annotation rankings between the two models (Figure 4B). We quantified the significance of this relationship using Pearson’s correlation, which revealed a moderately positive correlation (*r* = 0.3638, p-value = 3.52 *×* 10^−218^). For some V-J gene pairs, parameterizing microhomology has a particularly pronounced impact on sequence annotation. For example, sequences involving TRAV12-2*03 and TRAJ22*01 show a striking difference in annotation predictions between models, with 81.67% of sequences exhibiting different top-ranked annotations. This effect may be driven by the relatively high microhomology content across annotations for these sequences, averaging 0.1280 nucleotides compared to the overall average of 0.1144 nucleotides across all V-J gene pairs.

**Figure 4:**
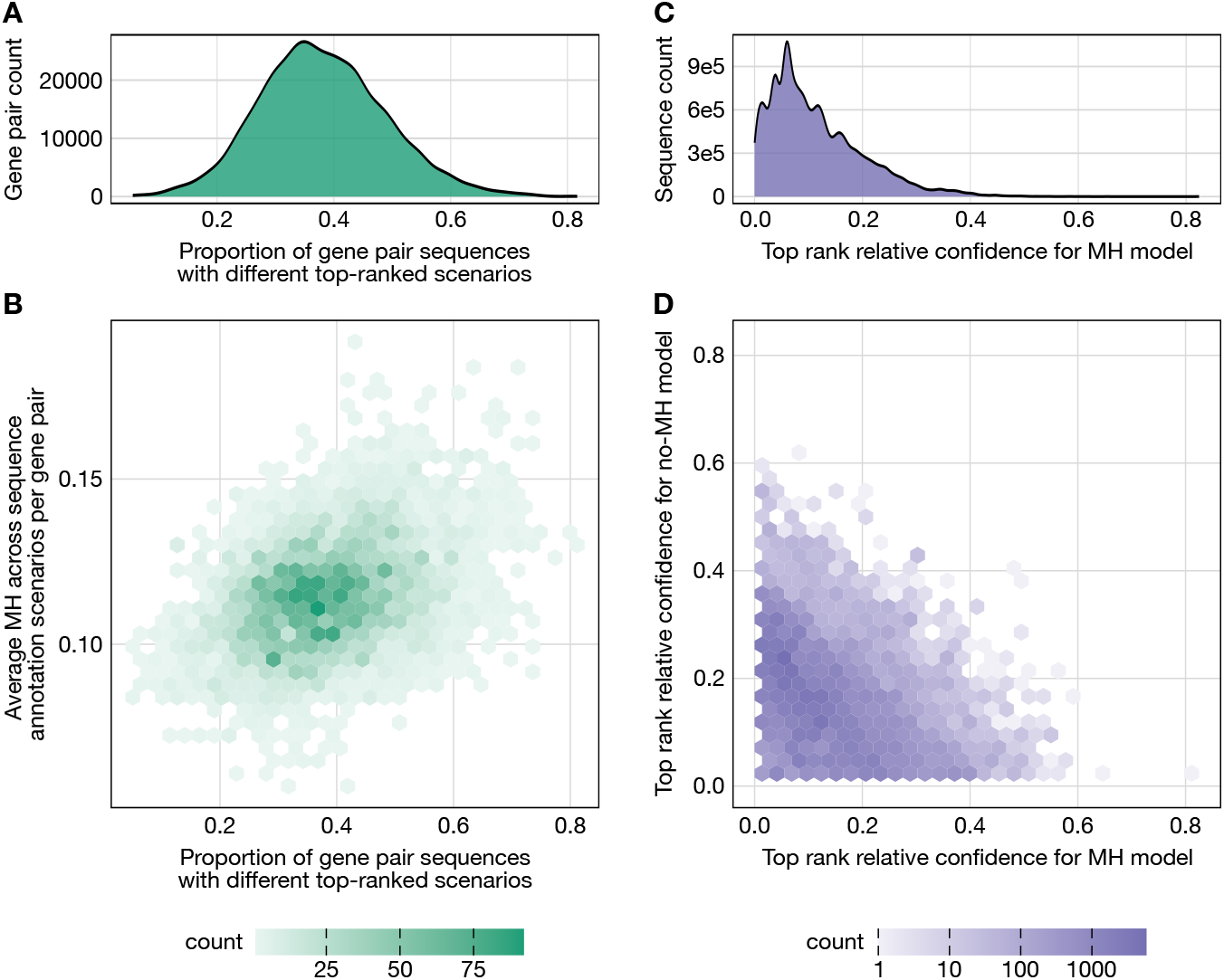
Accounting for microhomology (MH) in models used for sequence annotation substantially alters both V(D)J recombination annotation probabilities and rankings. **(A)** Distribution of the proportions of sequences per V-J gene pair that have a different top-ranked V(D)J recombination annotation when using the model that parameterizes microhomology to calculate annotation probabilities compared to using the model that does not. Only sequences with multiple possible annotations are considered when calculating each proportion. There is a correlation between the proportion of sequences with differing top-ranked annotations and the average microhomology across potential annotations per sequence across V-J gene pairs. This highlights how microhomology influences ranking changes across V-J gene pairs. **(C)** Distribution of the relative confidence for the top-ranked annotation using the MH model. Relative confidence is defined as the absolute difference in annotation probabilities using the MH model, comparing the top-ranked annotation from the MH model and the top-ranked annotation from the no-MH model for each sequence. **(D)** Comparison of the relative confidences of the top-ranked annotation for each sequence between the two models. As established in panel C, this involves comparing the absolute differences in annotation probabilities for the two top-ranked annotation of each sequence between the two models. Most sequences exhibit substantial shifts in relative annotation confidence between models, highlighting large model-driven changes in sequence annotations. These shifts occur even in cases where one or both models show high relative confidence.

Given that parameterizing microhomology leads to different top-ranked annotations for many sequences, we next quantified the relative confidence of these rankings. To explore this, we compared the annotation probabilities, as determined by the model with microhomology, between the top-ranked annotations from the models with and without microhomology. We define the relative confidence of the topranked annotation for the microhomology model as the absolute difference in these annotation probabilities. On average, for sequences containing a different top-ranked annotation between the two models, we find that the relative confidence of the top-ranked annotation for the microhomology model is 0.1140 (Figure 4C).

Additionally, we examined the relative confidence levels of the top-ranked annotations from both models. If microhomology merely resolved ties between competing annotations, we would expect minimal relative confidence in the top-ranked annotation using the model lacking microhomology terms, with larger relative confidence observed for the model containing microhomology terms. However, our findings indicate substantial shifts in relative confidence across models for most sequences (Figure 4D), with data points widely distributed rather than clustering near the axes. For instance, even when the model lacking microhomology terms has high confidence in its top-ranked annotation relative to the top-ranked annotation derived from the model containing microhomology, a similar flip in confidence is often observed when switching models. This effect suggests that parameterizing microhomology leads to meaningful changes in the annotation ranking landscape, potentially altering the biological interpretation of many sequences.

## Discussion

Previous *in vitro* experiments have suggested that microhomology plays a significant role in biasing key V(D)J recombination processing steps, such as trimming and ligation. However, these findings do not fully elucidate the importance of microhomology in shaping *in vivo* recombination outcomes in humans. In this paper, we use statistical inference on previously-published high-throughput human TCR repertoire data [23, 22] to assess whether microhomology influences V(D)J recombination in humans with intact recombination machinery. Our probabilistic modeling framework quantifies how sequence-level features, particularly microhomology, impact trimming and ligation decisions during V(D)J recombination. We find that (1) microhomology significantly increases trimming and ligation probabilities, (2) microhomology is an important predictor of recombination choices across multiple receptor loci and sequence types, and (3) accounting for microhomology when inferring V(D)J recombination annotations substantially changes annotation probabilities and rankings.

Our results reveal that microhomologous nucleotides between gene ends significantly increase ligation probabilities, aligning with previous *in vitro* evidence suggesting that microhomology guides ligation [14, 30, 28, 37, 8, 7, 38]. While much of the previous experimental focus has been on terminal microhomology (present at gene ends), many gene pairs lack terminal microhomology but have interior regions of microhomology. It has been proposed that trimming can expose these interior regions, which then guide ligation through microhomology-mediated processes [8, 7]. Our findings support this, as microhomology appears to have a stronger effect on trimming than on ligation, likely due to the dependence of ligation options on prior trimming choices. Because this analysis focuses on sequences without N-insertions—– allowing us to directly identify germline-microhomologymediated ligation events—–the observed strength of these effects may be amplified compared to analyses that include all sequences. Notably, when analyzing sequences with N-insertions—–where ligation is not mediated by germline microhomology—–we still observe that microhomology influences trimming, though less strongly, suggesting a more complex role for microhomology in V(D)J recombination beyond its involvement in ligation.

In addition to microhomology-related parameters, our modeling framework included sequence-level parameters designed to capture the effects of local nucleotide context, absolute trimming site positioning, and sequence breathing capacity. These parameters, except for the microhomology-related ones, were introduced in our previous analyses of trimming patterns for individual V- and J-gene sequences [41]. Our current results were consistent with those earlier findings. Specifically, parameters capturing the effects of absolute trimming site positioning and sequence breathing capacity downstream of the trimming site had the most substantial impact on improving overall model fit and predictive accuracy, showing the largest negative effect sizes on trimming and ligation choices. In contrast, microhomology-related parameters showed the largest positive effect sizes and consistently enhanced predictive accuracy. This pattern held when evaluating model performance and accuracy using sequences from different receptor loci and productivity types, underscoring microhomology’s role in recombination decisions.

Beyond its intrinsic interest, microhomology has significant practical implications. In addition to influencing trimming and ligation probabilities, we found that parameterizing microhomology-related effects leads to substantial shifts in V(D)J recombination annotation probabilities and rankings. These shifts often corresponded to large changes in the relative confidence of annotation rankings when comparing models that incorporate microhomology with those that do not. Such changes could significantly alter the annotation landscape, potentially impacting the biological interpretation of many sequences. Despite this, to our knowledge, all frequently used V(D)J recombination annotation software [45, 3, 32] do not account for microhomology or consider annotations that incorporate microhomologous nucleotides.

Our work has several limitations. First, we rely on non-productive rearrangements as a proxy for pre-selection recombination statistics, as is common in the literature [40, 33, 32, 44]. However, we acknowledge that these recombination statistics may not perfectly reflect the pre-selection repertoire. Second, our analysis excluded sequences with N-insertions, allowing us to use known germline V- and J-gene sequences to identify regions of germline-encoded microhomology and potential microhomology-mediated ligation events. N-insertions complicate the analysis because the identities of inserted nucleotides are unknown, and their presence suggests germline microhomology did not guide ligation. Because the presence or absence of N-insertions may affect the probability of successful ligation, excluding these sequences could shift the distribution of observed trimming and ligation events. Consequently, the microhomology effects that we have inferred may not extend to sequences with N-insertions. Future work could explore microhomology dynamics in sequences containing N-insertions, but doing so would require assumptions about latent variables such as N-insertion identities prior to ligation, making this analysis challenging if using repertoire sequencing data.

Despite the clear role of microhomology in biasing V(D)J recombination events and influencing V(D)J recombination annotation inference, no probabilistic models incorporating microhomology have been developed, to our knowledge. Future work could integrate microhomology-related dependencies into existing probabilistic frameworks like IGoR [32], which currently models dependencies between recombination events such as V- and J-gene choice, V-gene choice and V-gene deletions, and J-gene and J-gene deletions for TCR*α* sequences. To explicitly account for microhomology, additional dependencies would need to be introduced between V- and J-gene deletions, V-gene choice and J-gene deletion, and J-gene choice and V-gene deletion, along with incorporating new parameters to capture the sharing of microhomologous nucleotides. However, this approach could be challenging due to the large number of parameters required and the corresponding need for large datasets to adequately train the model. Alternative methods may need to be explored, especially to balance model complexity with practicality.

In summary, our findings demonstrate that microhomology plays a significant role in trimming and ligation choices during V(D)J recombination, underscoring the importance of accounting for microhomology effects when predicting recombination outcomes and annotating sequences. By advancing our understanding of microhomology’s influence in human V(D)J recombination, these results provide another step toward uncovering how this process generates diverse receptors that support a robust immune response in humans.

## Materials and Methods

### TCR*α* training dataset

We downloaded TCR*α* repertoire sequence data from thymocyte samples of 10 immunologically healthy infants (aged 0-1 years) from the Adaptive Biotechnologies immuneACCESS database, following links provided in the original publications [23, 22]. We assigned V(D)J recombination annotations to each sequence using the IGoR software (version 1.4.0) [32], which outputs potential recombination annotations along with their corresponding likelihoods. For each sequence, we inferred the ten highest probability recombination annotations and annotated each sequence by sampling from these annotations based on their posterior probabilities.

In preparing the training dataset, we implemented several filtering steps. First, we excluded sequences that had more than fourteen nucleotides trimmed from either the V-gene or J-gene, as more extensive trimming is uncommon and could suggest an alternative trimming mechanism. Next, we focused only on non-productive sequences to avoid the potential confounding effects of selection in recombination statistics. Additionally, we excluded sequences with N-insertions, as inferred by IGoR, to focus on potential germline-microhomology-mediated ligation events, since the presence of N-insertions indicates that ligation likely did not involve germline microhomology.

After applying these filtering criteria, we compiled a training dataset consisting of 1,257,528 sequences. Additionally, to validate our model, we used a separate dataset of 983,514 productive sequences.

### TCR*α* testing dataset

We downloaded TCR*α* repertoire sequence data from peripheral blood samples of 10 healthy individuals (aged 3-14 years) from the Adaptive Biotechnologies immuneACCESS database using the link in the original publication [22]. This cohort differs from the training cohort in demographics and sampling location. All individuals were heterozygous for HLA-DR3/DR4 and had siblings diagnosed with Type-1 Diabetes, but they showed no clinical symptoms of diabetes at the time of sampling or in the subsequent years. We applied the same IGoR-based annotation and filtering procedures to this testing dataset as used for the training dataset. For model validation, we separately analyzed 98,244 non-productive and 141,676 productive sequences.

### TCR*γ* testing dataset

Annotated TCR*γ* repertoire sequence data for 23 healthy bone marrow donor subjects was downloaded from the Adaptive Biotechnologies immuneACCESS database [39]. We applied the same filtering procedures to this testing dataset as used for the training dataset. For model validation, we separately analyzed approximately 44,673 non-productive and 20,681 productive sequences.

### Notation and modeling set-up

We explore the impact of microhomology on V(D)J recombination by modeling the joint probability of trimming and ligation scenarios given V-gene and J-gene sequences. To accurately assess the effects of microhomology, we focus our analysis on sequences without N-insertions. This approach allows us to leverage germline V- and J-gene sequences to identify potential germline-microhomology-mediated ligation events, as illustrated in Figure 1. We exclude sequences with N-insertions because their unknown pre-ligation nucleotide composition complicates the analysis of ligation dynamics and indicates a lack of *germline*-microhomology-mediated ligation, thus impeding our ability to quantify the influence of microhomology on the ligation process.

In order to set up our model, we will now summarize relevant notation. We uniformly sample a sequence, *X*, from a TCR*α* repertoire of filtered sequences, assuming each sequence is annotated with a specific V-gene, J-gene, number of nucleotides deleted from each gene, and number of N-insertions between the two genes. The following variables are random due to the choice of *X*, but are deterministic given *X*, as they are determined by sampling from the recombination annotations inferred by IGoR based on their posterior probabilities. Let *S*_*V*_ and *S*_*J*_ be random variables representing the V-gene and J-gene, respectively, and *I* be a random variable representing the number of N-insertions. Let *R*_*V*_ and *R*_*J*_ be random variables that represent the number of nucleotides deleted from each gene. For notational convenience we assume *R*_*V*_ and *R*_*J*_ each take on an integer value on the interval [−2, …, 14], where values outside this range are considered nonsensical and assigned a probability of zero. Negative values indicate P-nucleotide deletions: a trim of 0 means the deletion stops at the end of the germline gene sequence (e.g. two P-nucleotides are trimmed off), while a trim of −2 indicates no deletion of P-nucleotides or gene sequence nucleotides. This indexing is consistent with the IGoR software [32]. We define *R* and *S* as ordered pairs of IGoR-inferred trimming amounts and genes: *R* = (*R*_*V*_, *R*_*J*_) and *S* = (*S*_*V*_, *S*_*J*_).

Since existing sequence annotation methods like IGoR [32] do not consider microhomology and attribute shared nucleotides to only one sequence, we introduce a procedure for adjusting annotations to account for microhomology. We introduce *M* as a random variable representing the count of shared microhomologous nucleotides in the final observed sequence and *T*_*V*_ and *T*_*J*_ as random variables representing the microhomology-adapted trimming amounts. We define *T* = (*T*_*V*_, *T*_*J*_) as an ordered pair of microhomology-adapted trimming amounts, referring to *T* as a “trimming scenario” and *M* as a “ligation scenario.”

For each gene pair *S*, each IGoR-inferred trimming annotation *R* maps to a set of possible microhomology-adapted annotations described by *T* and *M*. While both *T* and *M* can be considered random variables, they are dependent on one another—meaning the possible values of *M* are constrained by *T* and vice versa. Using these mappings, we can form a set *A*_*X*_ that includes all feasible combinations of trimming and ligation scenarios that could explain the observed sequence *X* (as illustrated in Figure 1B). The process of obtaining *A*_*X*_ is explained in detail in the SI Appendix. We aim to model trimming and ligation scenario probabilities given V- and J-gene pairs. Let *Q* represent the sequence productivity of all sequences considered in our downstream modeling, which is deterministic and can be productive, nonproductive, or both. To estimate the empirical conditional probability density function, let *C*(*S, Q, I* = 0) represent the count of TCRs within a sampled repertoire with productivity *Q*, using gene pair *S*, and with zero N-insertions. Let *C*(*T, M, S, Q, I* = 0) represent the count with trimming scenario *T*, ligation scenario *M*, zero N-insertions, productivity *Q*, and gene pair *S*. The empirical conditional probability density function is defined as:

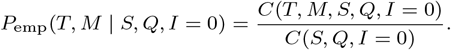

Ultimately, we will train a model of *P* (*T, M* | *S, Q, I* = 0) using sequence-level parameters, including those that capture microhomology-related effects, with our TCR*α* repertoire training dataset. However, since the true trimming and ligation annotations (*T, M*) for each sequence are latent variables, we cannot compute or estimate this probability density directly. To address this, we assign probabilities to each potential annotation based on their model likelihood. Because these probabilities depend on our model’s parameters, we use an expectation-maximization algorithm for parameter inference, which we describe in detail in subsequent sections. We summarize all the notation discussed in this section, as well as in the following sections, in Table S1.

### Model formulation and training

We employ a two-step conditional logit model to capture the decision-making process involved in selecting trimming and ligation scenarios for V-J gene pairs. Our model describes a generative process in two steps:

1. We model the probability, *P* (*T* | *S, Q, I* = 0), of choosing a trimming scenario *T* for a given V-J gene pair *S*, sequence productivity *Q*, and N-insertion amount *I* = 0. This probability is modeled by parameters specific to trimming scenarios.
2. We model the probability, *P* (*M* | *T, S, Q, I* = 0), of choosing a ligation scenario *M* for a given trimming scenario *T*, V-J gene pair *S*, sequence productivity *Q*, and N-insertion amount *I* = 0. This probability is modeled by parameters specific to each ligation scenario.

The joint probability of a trimming scenario *T* and a ligation scenario *M* for a given V-J gene pair *S*, sequence productivity *Q*, N-insertion amount *I* = 0 can be factored as:

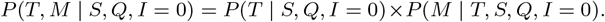

Figure 5 depicts the two-step structure of our model, illustrating the decision-making process for an example V-J gene pair.

**Figure 5:**
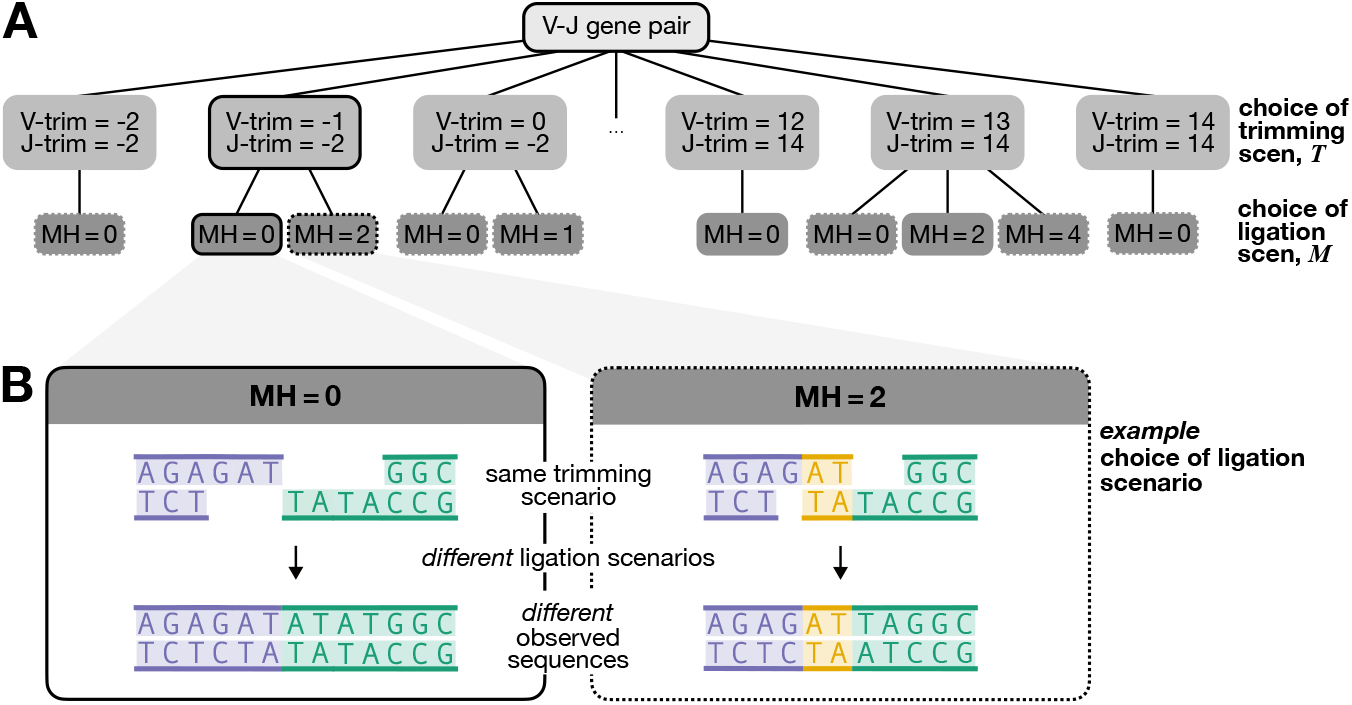
**(A)** Schematic of trimming and ligation choices for an arbitrary V-J gene pair, denoted by the random variable *S*. The first choice is the trimming scenario, represented by the random variable *T*, which consists of a pair of V- and J-gene trimming amounts (e.g. each can range from −2 to 14 nucleotides). The next choice is the ligation scenario, represented by the random variable *M*, which captures the number of microhomologous nucleotides used. The available ligation scenario choices depend on the germline sequences of the two genes being joined. Trimming and ligation scenarios resulting in productive and nonproductive sequences are shown in solid and dashed boxes, respectively. **(B)** Illustration of the possible ligation scenarios for an example pair of trimmed sequences. The chosen ligation scenario affects the resulting observed sequence. The trimmed V-gene sequence is shown in purple, the trimmed J-gene sequence in green, and microhomologous nucleotides in yellow. Deletions are indexed such that a deletion of 0 corresponds to the end of the germline gene sequence (two P-nucleotides trimmed) and −2 corresponds to the full sequence (no P-nucleotides trimmed). See Figure S8 for a version of this figure using variable notation.

To incorporate characteristics of each possible trimming and ligation scenario in our model, we define parameters specific to trimming and ligation scenarios. In our previous work, we established that local nucleotide identities at trimming sites (the “trimming motif”) and the counts of GC or AT nucleotides beyond these motifs (the “two-side base-count beyond”) are strong predictors of trimming probabilities for single gene sequences [41].Building on this foundation, we have integrated these established parameters with newly developed microhomology-related parameters to assess the combined effects on the processes of trimming and ligation.

First, we define a set of trimming-related parameters ***β***_trim_, which includes previously established trimming motif and base-count-beyond parameters for each gene, and a new trimming-related microhomology parameter. This new parameter measures the importance of the average number of microhomologous nucleotides between two trimmed sequences, a value that varies depending on the chosen trimming scenario. Similarly, we definea set of ligation-related parameters ***β***_lig_, which in our case only includes one parameter. This single parameter measures the importance of the number of microhomologous nucleotides between two trimmed and ligated sequences, which varies depending on the chosen trimming and ligation scenario. All of these parameters are summarized in Table S2 and defined in detail in the SI Appendix. With these parameters ***β***_trim_ and ***β***_lig_, our model estimates the joint probability of a trimming and ligation scenario (given by *T* and *M*) for a given V-J gene pair *S*, sequence productivity *Q*, and N-insertion amount *I* = 0:

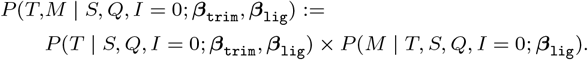

Training this model is complex because the true trimming and ligation annotation of each sampled sequence is a latent variable that will depend on the model parameters. As described earlier, we obtain the set of possible microhomology-adapted trimming and ligation annotations (*T* and *M*), denoted as *A*_*X*_, for a given sequence *X* by transforming the initial IGoR-inferred trimming annotations *R*. We assign probabilities (or weights) to each potential annotation, and since these probabilities depend on the model parameters, we use an expectation-maximization algorithm for parameter inference. An extended description of this model formulation and training details can be found in the SI Appendix.

### Assessing significance of model parameters

When training our model, we infer a set of model parameters 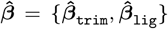 where ***β***_trim_ are trimming-related parameters and ***β***_lig_ are ligation-related parameters. To assess the significance of each parameter 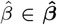, we test the null hypothesis that 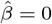. To do this, we estimate the standard error of each inferred parameter using a bootstrap method, with observed sequences as the sampling unit. For each bootstrap iteration, we sample sequences from the training dataset with replacement and train a new model to re-estimate the parameters. This process is repeated 1000 times, resulting in 1000 parameter estimates, which we use to calculate the standard error for each parameter. Using the estimated standard error, we test whether 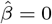 by calculating the test statistic:

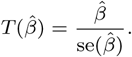

We compare 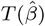 to a *N* (0, 1) distribution to obtain each p-value. We assess the significance of each model parameter using a Bonferroni-corrected threshold, adjusting for the total number of parameters being evaluated in the model.

### Validating model using likelihood ratio testing

To validate our model, we compare our model which contains microhomology-related terms to a relatively simpler model lacking these terms to determine which better fits the observed data. A likelihood ratio test (LRT) allows us to test the goodness-of-fit between these two nested models, where the complex model includes additional parameters (in this case, the microhomology-related parameters). The LRT statistic compares the log-likelihood scores of the two models:

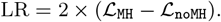

Here, *L*_MH_ represents the log-likelihood defined in the SI Appendix, (20). The log-likelihood score given by *L*_noMH_ is defined similarly, but the model lacks microhomology-related parameters. The LRT statistic approximately follows a chi-square distribution. The degrees of freedom for this test are equal to the number of additional parameters in the more complex model (e.g. 2 microhomology-related parameters in our case). Using this information, we calculate the p-value of the test statistic to determine whether the inclusion of microhomology-related parameters significantly improves the model fit. This helps us assess whether these parameters are biologically meaningful given our observed data.

## Data and code availability

Code implementing the modeling described is available at https://github.com/magdalenarussell/microhomology. All models and optimization algorithms are implemented using the JAX and JAXopt packages, which support automatic differentiation [5, 2]. The data used in this study were previously published and can be accessed through the Adaptive Biotechnologies immuneACCESS database via the links provided in the original publications [39, 23, 22].

## Acknowledgements

The authors thank Armita Nourmohammad and Michael Lieber for helpful discussions regarding this paper. The authors also acknowledge Fred Hutch Scientific Computing, supported by the National Institutes of Health (NIH) award S10OD028685. This work was supported by the NIH through awards R01 AI146028 (M.L.R., P.B., F.A.M.), R01 AI136514 (P.B.), R35 GM142795 (A.T.), and R35 GM141457 (P.B.). Additional support for A.T. was provided by the Mahan Fellowship. Dr. Matsen is an Investigator of the Howard Hughes Medical Institute (HHMI). This article is subject to HHMI’s Open Access to Publications policy. HHMI lab heads have previously granted a nonexclusive CC BY 4.0 license to the public and a sublicensable license to HHMI in their research articles. Pursuant to those licenses, the author-accepted manuscript of this article can be made freely available under a CC BY 4.0 license immediately upon publication.

## Footnotes

1 In this work, we use the term “terminal microhomology” to describe *germline-encoded* microhomologous nucleotides located at gene ends. However, other sources [28, 37] often use the term more broadly to describeall microhomologous nucleotides located at gene ends, including both germline-encoded nucleotides and those generated through N-insertion.

