## Supplemental Materials for "Statistical analysis of repertoire data demonstrates the influence of microhomology in V(D)J recombination"

### Contents

**1 Supporting Information Text** **1**

**2 Supporting Figures and Tables** **11**

**3 SI References** **22**

### 1 Supporting Information Text

#### Identifying the set of possible annotations for a sequence

We aim to identify all feasible combinations of microhomology-adapted trimming and ligation values ($T_V$, $T_J$, and $M$) that could account for the observed sequence $X$. Recall that $S_V$ and $S_J$ represent the V-gene and J-gene, which each define a V-gene and J-gene sequence, respectively. For ease of notation, these sequences are both oriented in the 3'-to-5' direction and are represented as ordered lists of nucleotides. Although the top strand of the V-gene typically follows a 5'-to-3' orientation, we reverse it here for notational convenience.

To begin, we employ a sequence annotation tool such as IGoR to infer a trimming configuration, given by $R_V$ and $R_J$, denoted as $r_V$ and $r_J$ for specific initial instances. Because these annotation tools do not consider microhomology, they assume that $M = 0$. Thus, our initial set of possible annotations, $A_X$, includes only $(T_V = r_V, T_J = r_J, M = 0)$. To expand $A_X$ to include annotations with microhomology $(M > 0)$, we introduce a function $h(x, y)$, which quantifies the number of contiguous complementary nucleotides between two overlapping, equal-length sequence regions $x$ and $y$. This function is defined as:

$$h(x, y) = \sum_{i=0}^{\text{len}(x)} \begin{cases} 1 & \text{if } x(j) \text{ is complementary to } y(j) \text{ for all } j \in \{0, \dots, i\} \\ 0 & \text{otherwise.} \end{cases} \tag{1}$$

We apply this function to assess complementarity in overlapping regions between a V-gene sequence $S_V$ and a J-gene sequence $S_J$, aligning the sequences without gaps at the IGoR-inferred trimming sites $r_V$ and $r_J$ (Figure S9). The overlapping regions are:

1. The *V-gene:trimmed-J* region, where $\texttt{seq}_{\texttt{trimmed}}(S_J, r_J)$ represents the trimmed J-gene sequence oriented 5'-to-3', and $\texttt{seq}_{\texttt{overlap}}(S_V, r_V, r_J)$ represents the overlapping V-gene sequence oriented 3'-to-5'.

2. The *J-gene:trimmed-V* region, with $\texttt{seq}_{\texttt{trimmed}}(S_V, r_V)$ and $\texttt{seq}_{\texttt{overlap}}(S_J, r_J, r_V)$ representing the trimmed V-gene and overlapping J-gene sequences oriented 5'-to-3' and 3'-to-5', respectively.

We define $k_J$ and $k_V$ as the counts of contiguous complementary nucleotides in these regions:

$$k_J = h\big(\texttt{seq}_{\texttt{overlap}}(S_V, r_V, r_J), \texttt{seq}_{\texttt{trimmed}}(S_J, r_J)\big)$$

and

$$k_V = h\big(\mathsf{seq}_{\texttt{overlap}}(S_J, r_J, r_V), \mathsf{seq}_{\texttt{trimmed}}(S_V, r_V)\big).$$

Given these values, a sequence $X$ with an initial IGoR-inferred trimming configuration ($T_V = r_V, T_J = r_J, M = 0$) can also be annotated with microhomologous nucleotide counts $M$ ranging from 0 to $k_V + k_J$. For each of these values of $M$, the corresponding trimming amounts ($T_V$ and $T_J$) are adjusted accordingly. This expands the set of possible annotations $A_X$ to:

$$A_X = \{(r_V, r_J, 0)\} \cup \{(r_V, r_J, m) \mid \text{conditions}\}.$$

The conditions are:

1. For each possible annotation, $t_V$ and $t_J$ (realizations of $T_V$ and $T_J$) are within the range of initial IGoR-inferred trimming values adjusted by the contiguous nucleotide count: $r_V - k_V \leq t_V \leq r_V$ and $r_J - k_J \leq t_J \leq r_J$.

2. The microhomologous nucleotide count $m$ (a realization of $M$) ranges from 0 to the sum of contiguous complementary nucleotides: $0 \leq m \leq k_V + k_J$.

3. The sum of the trimming amounts $t_V$ and $t_J$ and the microhomologous nucleotide count $m$ equals the sum of the initial IGoR-inferred trimming amounts: $t_V + t_J + m = r_V + r_J$

This process can be repeated to identify the sets of all possible sequence annotations for each sequence sampled from a TCR$\alpha$ repertoire.

#### Defining a model weight function

We aim to model the influence of various sequence-level parameters, including microhomology-related parameters, on joint trimming and ligation configuration probabilities, $P(T, M \mid S, Q, I = 0)$, where $T$ represents the microhomology-adapted trimming configuration, $M$ represents the number of microhomologous nucleotides used in ligation, $S$ represents the gene pair, $Q$ represents the productivity of the sequences, and $I = 0$ represents zero N-insertions. For our modeling purposes, we assume the following about V(D)J recombination biology:

1. The DNA hairpin of each joining gene is nicked open by a single-stranded break [3, 8, 7, 4, 6].

2. This hairpin nick occurs at the +2 position, creating a 4-nucleotide-long 3'-single-stranded overhang, with the two 3'-most nucleotides being P-nucleotides [7, 6].

3. If any part of the original gene sequence is deleted, all P-nucleotides will also be deleted [3, 10].

These assumptions allow us to determine the germline nucleotide sequence on both sides of each trimming site and define sequence-level model features. We assume that observations can be drawn from a model where these features vary across trimming and/or ligation configurations for a given gene pair. Using these assumptions, we previously demonstrated that local nucleotide identity surrounding trimming sites (the "trimming motif") and the counts of GC or AT nucleotides beyond these motifs (the "two-side base-count beyond") are highly predictive of trimming probabilities for single gene sequences [9]. Building on this foundation, we aim to integrate these established parameters with newly developed microhomology-related parameters to assess the combined effects on the processes of trimming and ligation.

For our model, we define two sets of parameters, one trimming-related and one ligation-related, to model the probabilities of trimming and ligation configurations. We model trimming configuration probabilities using established trimming motif (given by $\boldsymbol{\beta}_V^{\texttt{motif}}$ and $\boldsymbol{\beta}_J^{\texttt{motif}}$) and two-side base-count beyond (given by $\boldsymbol{\beta}_V^{\texttt{AT}}$, $\boldsymbol{\beta}_J^{\texttt{AT}}$, $\boldsymbol{\beta}_V^{\texttt{GC}}$, and $\boldsymbol{\beta}_J^{\texttt{GC}}$) parameters for each gene, along with a new parameter related to microhomology (given by $\beta^{\texttt{trimMH}}$). This new parameter measures the importance of the average number of microhomologous nucleotides between two trimmed sequences for trimming configuration probabilities. Additionally, we model ligation configuration probabilities using another new parameter related to microhomology (given by $\beta^{\texttt{ligMH}}$), which measures the importance of the number of microhomologous nucleotides within the ligation configuration.

Using these model parameters, we define weight functions for the trimming choice $f_{\texttt{trim}}$ and the ligation choice $f_{\texttt{lig}}$ such that our model of $P(T, M \mid S, Q, I = 0)$ will be a normalized version of these weights.

The trimming-related weight function $f_{\texttt{trim}}$ aggregates the desired parameter-specific weight functions (defined in Table S2) as follows:

$$\begin{aligned} &f_{\texttt{trim}}(T, S; \boldsymbol{\beta}_{\texttt{trim}}) \\ &\qquad := f_{\texttt{motif}}(T_V, S_V; \boldsymbol{\beta}_V^{\texttt{motif}}) + f_{\texttt{motif}}(T_J, S_J; \boldsymbol{\beta}_J^{\texttt{motif}}) \\ &\qquad + f_{\texttt{count}}(T_V, S_V; \boldsymbol{\beta}_V^{\texttt{AT}}, \boldsymbol{\beta}_V^{\texttt{GC}}) + f_{\texttt{count}}(T_J, S_J; \boldsymbol{\beta}_J^{\texttt{AT}}, \boldsymbol{\beta}_J^{\texttt{GC}}) \\ &\qquad + f_{\texttt{trimMH}}(T, S; \beta^{\texttt{trimMH}}). \end{aligned} \tag{2}$$

For notational convenience, $\boldsymbol{\beta}_{\texttt{trim}}$ represents the set of all trimming-related regression parameters, $S = (S_V, S_J)$ represents a gene pair, $T = (T_V, T_J)$ represents a trimming configuration, and $M$ represents the number of microhomologous nucleotides within the ligation configuration.

Additionally, we define a ligation-related weight function $f_{\texttt{lig}}$ that consists of the relevant parameter-specific weight function (defined in Table S2) as follows:

$$f_{\texttt{lig}}(T, M, S; \boldsymbol{\beta}_{\texttt{lig}}) := f_{\texttt{ligMH}}(M; \beta^{\texttt{ligMH}}) \tag{3}$$

where $\boldsymbol{\beta}_{\texttt{lig}}$ represents the set of all ligation-related regression parameters, which for our purposes includes only $\beta^{\texttt{ligMH}}$.

We further define each of these trimming-related and ligation-related regression parameters, along with their corresponding weight functions, within the following sections:

##### Defining "trimming motif" parameters

As in our previous work [9], we define trimming motif parameters to include one nucleotide position 5' of the trimming site and two nucleotide positions 3' of the trimming site. We describe this definition for a V-gene sequence and V-gene trimming amount, but it applies similarly to a J-gene sequence and J-gene trimming amount.

Recall that $S_V$ represents the V-gene which defines a V-gene sequence. For ease of notation, we orient this sequence in the 3'-to-5' direction and represent it as an ordered list of nucleotides. Recall that $R_V$ is a random variable representing a V-gene trimming amount. Let $S_V(R_V + 2 - j)$ represent the nucleotide identity at the trimming motif position $j \in \{0, \ldots, 2\}$ where positions $j \leq 0$ represent motif positions 5' of the trimming site and positions $j > 0$ represent motif positions 3' of the trimming site. The trimming motif sequence (oriented 5'-to-3') is given by the ordered list:

$$\big(S_V(R_V + 2 - j)\big)_{j=0}^{2}. \tag{4}$$

Depending on $R_V$, this trimming motif may or may not include P-nucleotides. For $R_V \geq 2$, the two 3' trimming motif nucleotides will include the two deleted gene sequence nucleotides 3' of the trimming site (and no P-nucleotides). Since we are assuming that the initial hairpin nick occurs at the +2 position, there will be two P-nucleotides present in the 5'-to-3' gene sequence. For $0 \leq R_V < 2$, P-nucleotides will be included in the trimming motif sequence. Likewise, as a result of the +2 hairpin nick position assumption, TCRs that have $R_V < 0$ will not have a full-length nucleotide trimming motif. For these "off-the-end" motif cases, we assign zero influence to the missing nucleotides during model fitting.

Let $\beta_{jk}^{\texttt{motif}}$ be a (log) position-weight-matrix parameter for trimming motif position $j \in \{0, \ldots, 2\}$ and nucleotide $k \in \{A, T, C, G\}$. The set of all such parameters for the V-gene is denoted by $\boldsymbol{\beta}_V^{\texttt{motif}}$. We can define an un-normalized position-weight-matrix weight:

$$f_{\texttt{motif}}(R_V, S_V; \boldsymbol{\beta}_V^{\texttt{motif}}) := \sum_{j=0}^{2} \beta_{jS_V(R_V+2-j)}^{\texttt{motif}} \tag{5}$$

that will serve as a *motif*-specific weight function in subsequent modeling. As described above, since we are considering "off-the-end" motif cases, $S_V(R_V + 2 - j)$ will represent the nucleotide identity at sequence position $j$ where positions $j \leq 0$ represent sequence positions 5' of the trimming site and positions $j > 0$ represent sequence positions 3' of the trimming site.

##### Defining "base count" parameters

As in our previous work [9], we will also define parameters for the counts of GC and AT nucleotides on either side of each trimming site. We describe this definition for a V-gene sequence and V-gene trimming amount, but it applies similarly to a J-gene sequence and J-gene trimming amount. For an arbitrary sequence $x$, we can count the number of AT and GC nucleotides within the sequence as

$$C^{\texttt{AT}}(x) = C^{\texttt{A}}(x) + C^{\texttt{T}}(x) \tag{6}$$

and

$$C^{\texttt{GC}}(x) = C^{\texttt{G}}(x) + C^{\texttt{C}}(x), \tag{7}$$

respectively.

Since the count of AT or GC nucleotides within the sequences 5' and 3' of the trimming site may influence the probability of trimming differently, we calculate the counts separately and exclude nucleotides already included in the *motif* parameterization. As above, recall that $R_V$ represents a V-gene trimming amount and $S_V$ represents a V-gene which defines a V-gene sequence. For ease of notation, we orient this sequence in the 3'-to-5' direction and represent it as an ordered list of nucleotides. Let $S_V(R_V + 2 - j)$ represent the nucleotide identity at the trimming motif position $j \in \{0, \ldots, 2\}$ where positions $j \leq 0$ represent motif positions 5' of the trimming site and positions $j > 0$ represent motif positions 3' of the trimming site. As in our previous work, we include the ten nucleotides 5' of the motif in the 5' nucleotide counts. Since we include one nucleotide 5' of the trimming site in the "trimming motif" parameters (as described in the previous section), the nucleotide sequence 5' of the trimming site, beyond the "trimming motif", is given by the ordered list

$$\mathsf{seq}_5(R_V, S_V) = \big(S_V(R_V + 2 - j)\big)_{j=-11}^{-1}. \tag{8}$$

To count the number of AT and GC nucleotides in the sequence 3' of the trimming site, we include all nucleotides located 3' of the trimming site beyond the "trimming motif." Since we are interested in using GC nucleotide content as a proxy for sequence-breathing, which is relevant only for paired nucleotides, we exclude nucleotides within the 3' single-stranded overhang. Assuming the initial hairpin nick occurs at the +2 position, leading to a 4-nucleotide-long 3' single-stranded overhang, for $R_V > 2$, the nucleotide sequence 3' of the trimming site, beyond the "trimming motif" (which contains two nucleotide positions 3' of the trimming site), is given by the ordered list:

$$\mathsf{seq}_3(R_V, S_V) = \begin{cases} \big(S_V(R_V + 2 - j)\big)_{j=3}^{(R_V-2)} & \text{if } (R_V - 2) \geq 3 \\ () & \text{if } (R_V - 2) < 3. \end{cases} \tag{9}$$

For $(R_V - 2) < 3$, all nucleotides 3' of the trimming site are considered single-stranded, and thus no nucleotides will be included in the sequence used to calculate the AT and GC base-counts.

With these sequences 5' and 3' of the trimming site, we define $\beta^{\texttt{AT}}_{5V}$, $\beta^{\texttt{AT}}_{3V}$, $\beta^{\texttt{GC}}_{5V}$, and $\beta^{\texttt{GC}}_{3V}$ to be V-gene specific *base-count-beyond* model parameters for 5' and 3' sequence base-counts of AT and GC beyond the "trimming motif", respectively. The set of all such parameters for the V-gene are denoted by $\boldsymbol{\beta}^{\texttt{AT}}_V$ and $\boldsymbol{\beta}^{\texttt{GC}}_V$. With these parameters, we define a *base-count-beyond* weight function:

$$\begin{aligned} f_{\texttt{count}}(R_V, S_V; \boldsymbol{\beta}^{\texttt{AT}}_V, \boldsymbol{\beta}^{\texttt{GC}}_V) := \beta^{\texttt{AT}}_{5V} \cdot C^{\texttt{AT}}(\mathsf{seq}_5(R_V, S_V)) + \beta^{\texttt{AT}}_{3V} \cdot C^{\texttt{AT}}(\mathsf{seq}_3(R_V, S_V)) \\ + \beta^{\texttt{GC}}_{5V} \cdot C^{\texttt{GC}}(\mathsf{seq}_5(R_V, S_V)) + \beta^{\texttt{GC}}_{3V} \cdot C^{\texttt{GC}}(\mathsf{seq}_3(R_V, S_V)). \end{aligned} \tag{10}$$

using the functions $C^{\texttt{AT}}$ and $C^{\texttt{GC}}$ as defined in (6) and (7), respectively. As defined, these GC and AT base-counts for the 3' sequence are dependent on sequence length and provide a parameterization of both GC nucleotide content and length.

##### Defining "microhomology" parameters for trimming configuration choice

We can parameterize the average number of microhomologous nucleotides across possible ligation configuration choices for a given trimming configuration and define $\beta^{\texttt{trimMH}}$ to be an *microhomology* model parameter specific to trimming choice. We define a function $g$ that returns this average value as follows:

$$g(T, S) := \frac{\sum_{M' \in \mathcal{M}_{ST}} M'}{|\mathcal{M}_{ST}|}$$

such that $\mathcal{M}_{ST}$ is the set of all possible ligation configurations for the chosen trimming configuration $T$ and gene pair $S$. With this function, we can define a *microhomology* weight function

$$f_{\texttt{trimMH}}(T, S; \beta^{\texttt{trimMH}}) := \beta^{\texttt{trimMH}} \cdot g(T, S). \tag{11}$$

**Defining "microhomology" parameters for ligation configuration choice**

We can directly use the ligation configuration $M$, which represents the number of microhomologous nucleotides in the observed sequence, as a parameter. We then define $\beta^{\texttt{ligMH}}$ as the *microhomology* model parameter for predicting the choice of ligation configuration. We use the term "observed" for this microhomology parameter because these particular microhomologous nucleotides directly participate in the ligation process and are homologous in the final sequence. As such, we can define a *microhomology* weight function:

$$f_{\texttt{ligMH}}(M; \beta^{\texttt{ligMH}}) := \beta^{\texttt{ligMH}} \cdot M. \tag{12}$$

#### Extended model formulation and training description

We aim to model the influence of various sequence-level parameters, including microhomology-related parameters, on joint trimming and ligation configuration probabilities, $P(T, M \mid S, Q, I = 0)$, where $T$ represents the microhomology-adapted trimming configuration, $M$ represents the number of microhomologous nucleotides used in ligation, $S$ represents the gene pair, $Q$ represents the productivity of the sequences, and $I = 0$ represents zero N-insertions. Modeling this probability is complex because the true trimming and ligation annotation of each sampled sequence is a latent variable that will depend on the model parameters. As described earlier, we obtain the set of possible microhomology-adapted trimming and ligation annotations ($T$ and $M$), denoted as $A_X$, for a given sequence $X$ by transforming the initial IGoR-inferred trimming annotations $R$. We assign probabilities (or weights) to each potential annotation, and since these probabilities depend on the model parameters, we use an expectation-maximization algorithm for parameter inference. Below, we provide a detailed description of these steps.

We employ a two-step conditional logit model to capture the decision-making involved in selecting trimming and ligation configurations for V-J gene pairs. Our model describes a generative process in two steps:

1. We model the probability, $P(N \mid S, Q)$, of choosing a trimming configuration $N$ for a given V-J gene pair $S$ and sequence productivity $Q$. This probability is modeled by parameters specific to trimming configurations.

2. We model the probability, $P(M \mid N, S, Q)$, of choosing a ligation configuration $M$ for a given trimming configuration $N$, V-J gene pair $S$, and sequence productivity $Q$. This probability is modeled by parameters specific to each ligation configuration.

The joint probability of a trimming configuration $N$ and a ligation configuration $M$ for a given V-J gene pair $S$ and sequence productivity $Q$ can be factored as:

$$P(N, M \mid S, Q) = P(N \mid S, Q) \times P(M \mid N, S, Q).$$

Figure 3 depicts the two-step structure of our model, illustrating the decision-making process for an example V-J gene pair.

To incorporate characteristics of each possible trimming and ligation configuration in our model, we define parameter-specific weight functions such that our model of $P(T, M \mid S, Q, I = 0)$ will be a normalized version of these weights. In our previous work, we established that local nucleotide identities at trimming sites (the "trimming motif") and the counts of GC or AT nucleotides beyond these motifs (the "two-side base-count beyond") are strong predictors of trimming probabilities for single gene sequences [9]. Building on this foundation, we have integrated these established parameters with newly developed microhomology-related parameters to assess the combined effects on the processes of trimming and ligation. First, we define the trimming-related weight function $f_{\texttt{trim}}(T, S; \boldsymbol{\beta}_{\texttt{trim}})$ parameterized by as set of trimming-related parameters $\boldsymbol{\beta}_{\texttt{trim}}$, which includes previously established trimming motif and base-count-beyond parameters, and a new trimming-related microhomology parameter. This new parameter measures the effect of the average number of microhomologous nucleotides between two sequences, a value that varies depending on the chosen trimming configuration. Similarly, we define the ligation-related weight function $f_{\texttt{lig}}(T, M, S; \boldsymbol{\beta}_{\texttt{lig}})$ parameterized by a new ligation-related microhomology parameter $\boldsymbol{\beta}_{\texttt{lig}}$ which measures the effect of the number of microhomologous nucleotides between two sequences, a value that varies depending on the chosen trimming and ligation configuration. These parameters and weight functions are summarized in Table S2 and defined in detail in previous Supplementary Materials sections.

With these weight functions, our model estimates the joint probability of a trimming and ligation configuration (given by $T$ and $M$) for a given V-J gene pair $S$, sequence productivity $Q$, and N-insertion amount $I = 0$, combining influences of regression parameters $\boldsymbol{\beta}_{\texttt{trim}}$ and $\boldsymbol{\beta}_{\texttt{lig}}$:

$$P(T, M \mid S, Q, I = 0; \boldsymbol{\beta}_{\texttt{trim}}, \boldsymbol{\beta}_{\texttt{lig}}) := P(T \mid S, Q, I = 0; \boldsymbol{\beta}_{\texttt{trim}}, \boldsymbol{\beta}_{\texttt{lig}}) \times P(M \mid T, S, Q, I = 0; \boldsymbol{\beta}_{\texttt{lig}}). \quad (13)$$

To model the trimming configuration probability $P(T \mid S, Q, I = 0; \boldsymbol{\beta}_{\texttt{trim}}, \boldsymbol{\beta}_{\texttt{lig}})$, we expand conditional probability, giving:

$$P(T, S, Q, I = 0; \boldsymbol{\beta}_{\texttt{trim}}, \boldsymbol{\beta}_{\texttt{lig}}) = P(Q, I = 0 \mid T, S; \boldsymbol{\beta}_{\texttt{lig}}) \times P(T, S; \boldsymbol{\beta}_{\texttt{trim}}).$$

With this, we model $P(T \mid S, Q, I = 0; \boldsymbol{\beta}_{\texttt{trim}}, \boldsymbol{\beta}_{\texttt{lig}})$ as:

$$\begin{aligned} P(T \mid S, Q, I = 0; \boldsymbol{\beta}_{\texttt{trim}}, \boldsymbol{\beta}_{\texttt{lig}}) &= \frac{P(T, S, Q, I = 0; \boldsymbol{\beta}_{\texttt{trim}}, \boldsymbol{\beta}_{\texttt{lig}})}{\sum_{T' \in \mathcal{T}} P(T', S, Q, I = 0; \boldsymbol{\beta}_{\texttt{trim}}, \boldsymbol{\beta}_{\texttt{lig}})} \\ &= \frac{P(Q, I = 0 \mid T, S; \boldsymbol{\beta}_{\texttt{lig}}) \cdot P(T, S; \boldsymbol{\beta}_{\texttt{trim}})}{\sum_{T' \in \mathcal{T}} P(Q, I = 0 \mid T', S; \boldsymbol{\beta}_{\texttt{lig}}) \cdot P(T', S; \boldsymbol{\beta}_{\texttt{trim}})} \\ &:= \frac{P(Q, I = 0 \mid T, S; \boldsymbol{\beta}_{\texttt{lig}}) \cdot \exp\left(f_{\texttt{trim}}(T, S; \boldsymbol{\beta}_{\texttt{trim}})\right)}{\sum_{T' \in \mathcal{T}} P(Q, I = 0 \mid T', S; \boldsymbol{\beta}_{\texttt{lig}}) \cdot \exp\left(f_{\texttt{trim}}(T', S; \boldsymbol{\beta}_{\texttt{trim}})\right)} \end{aligned} \quad (14)$$

where $f_{\texttt{trim}}$ is the trimming-related weight defined in (2) and $\mathcal{T}$ is the set of all possible trimming configurations for the specified sequence productivity $Q$ and N-insertion amount $I = 0$. We model $P(Q, I = 0 \mid T, S; \boldsymbol{\beta}_{\texttt{lig}})$ as:

$$\begin{aligned} P(Q, I = 0 | T, S; \boldsymbol{\beta}_{\texttt{lig}}) = \frac{P(T, S, Q, I = 0; \boldsymbol{\beta}_{\texttt{lig}})}{P(T, S; \boldsymbol{\beta}_{\texttt{lig}})} &= \frac{\sum_{M_1 \in \mathcal{M}_1} P(M_1, T, S, Q, I = 0; \boldsymbol{\beta}_{\texttt{lig}})}{\sum_{M_2 \in \mathcal{M}_2} P(M_2, T, S; \boldsymbol{\beta}_{\texttt{lig}})} \\ &= \frac{\sum_{M_1 \in \mathcal{M}_1} P(M_1, T, S; \boldsymbol{\beta}_{\texttt{lig}})}{\sum_{M_2 \in \mathcal{M}_2} P(M_2, T, S; \boldsymbol{\beta}_{\texttt{lig}})} \\ &:= \frac{\sum_{M_1 \in \mathcal{M}_1} \exp\left(f_{\texttt{lig}}(T, M_1, S; \boldsymbol{\beta}_{\texttt{lig}})\right)}{\sum_{M_2 \in \mathcal{M}_2} \exp\left(f_{\texttt{lig}}(T, M_2, S; \boldsymbol{\beta}_{\texttt{lig}})\right)} \end{aligned} \quad (15)$$

given that $P(M, T, S, Q, I = 0; \boldsymbol{\beta}_{\texttt{lig}}) = P(M, T, S; \boldsymbol{\beta}_{\texttt{lig}})$. Here, $f_{\texttt{lig}}$ is the ligation-related weight defined in (12), $\mathcal{M}_1$ is the set of all possible ligation configurations for the chosen trimming configuration $T$, sequence productivity $Q$, and N-insertion amount $I = 0$ and $\mathcal{M}_2$ is the set of all possible ligation configurations for the chosen trimming configuration $T$. Similarly, we model the ligation configuration probability $P(M \mid T, S, Q, I = 0; \boldsymbol{\beta}_{\texttt{lig}})$ as:

$$\begin{aligned} P(M \mid T, S, Q, I = 0; \boldsymbol{\beta}_{\texttt{lig}}) &= \frac{P(M, T, S, Q, I = 0; \boldsymbol{\beta}_{\texttt{lig}})}{\sum_{M_1 \in \mathcal{M}_1} P(M_1, T, S, Q, I = 0; \boldsymbol{\beta}_{\texttt{lig}})} \\ &= \frac{P(M, T, S; \boldsymbol{\beta}_{\texttt{lig}})}{\sum_{M_1 \in \mathcal{M}_1} P(M_1, T, S; \boldsymbol{\beta}_{\texttt{lig}})} \\ &:= \frac{\exp\left(f_{\texttt{lig}}(T, M, S; \boldsymbol{\beta}_{\texttt{lig}})\right)}{\sum_{M_1 \in \mathcal{M}_1} \exp\left(f_{\texttt{lig}}(T, M_1, S; \boldsymbol{\beta}_{\texttt{lig}})\right)}. \end{aligned} \quad (16)$$

Combining these, our model becomes

$$\begin{aligned}
&P(T, M \mid S, Q, I=0; \boldsymbol{\beta}_{\texttt{lig}}, \boldsymbol{\beta}_{\texttt{trim}}) \\
&\qquad := P(T \mid S, Q, I=0; \boldsymbol{\beta}_{\texttt{trim}}, \boldsymbol{\beta}_{\texttt{lig}}) \times P(M \mid T, S, Q, I=0; \boldsymbol{\beta}_{\texttt{lig}}) \\
&\qquad := \frac{P(Q, I=0 \mid T, S; \boldsymbol{\beta}_{\texttt{lig}}) \cdot \exp\left(f_{\texttt{trim}}(T, S; \boldsymbol{\beta}_{\texttt{trim}})\right)}{\sum_{T' \in \mathcal{T}} P(Q, I=0 \mid T', S; \boldsymbol{\beta}_{\texttt{lig}}) \cdot \exp\left(f_{\texttt{trim}}(T', S; \boldsymbol{\beta}_{\texttt{trim}})\right)} \times \frac{\exp\left(f_{\texttt{lig}}(T, M, S; \boldsymbol{\beta}_{\texttt{lig}})\right)}{\sum_{M_1 \in \mathcal{M}_1} \exp\left(g(T, M_1, S; \boldsymbol{\beta}_{\texttt{lig}})\right)}.
\end{aligned} \tag{17}$$

where $P(Q, I=0 \mid T, S; \boldsymbol{\beta}_{\texttt{lig}})$ is defined in (15).

Recall that our data consists of sequences and we are considering sets of all possible sequence annotations $A_X$ for each sampled sequence $X$. Each annotation includes a microhomology-adapted trimming configuration $T$ ligation configuration $M$. To infer the parameters of our model $P(T, M \mid S, Q, I=0; \boldsymbol{\beta}_{\texttt{lig}}, \boldsymbol{\beta}_{\texttt{trim}})$, defined in 17, while marginalizing over all possible sequence annotations for each sequence, we employ an expectation-maximization (EM) approach. This iterative algorithm proceeds as follows: starting with initial model parameters $\boldsymbol{\beta}_{\texttt{lig}}$ and $\boldsymbol{\beta}_{\texttt{trim}}$, we aim to update to improved parameters $\boldsymbol{\beta}'_{\texttt{lig}}$ and $\boldsymbol{\beta}'_{\texttt{trim}}$. We define the normalized conditional probability of a specific sequence annotation $(T=t, M=m) \in A_X$ given a sequence $X$ with gene pair $S=s$ as:

$$P_{\text{annot}}(T=t, M=m \mid S=s, Q=q, I=0; X, \boldsymbol{\beta}_{\texttt{lig}}, \boldsymbol{\beta}_{\texttt{trim}}) = \frac{P(t, m \mid s, q, I=0; \boldsymbol{\beta}_{\texttt{lig}}, \boldsymbol{\beta}_{\texttt{trim}})}{\sum_{(t', m') \in A_X} P(t', m' \mid s, q, I=0; \boldsymbol{\beta}_{\texttt{lig}}, \boldsymbol{\beta}_{\texttt{trim}})}. \tag{18}$$

Here, $P(t, m \mid s, q, I=0; \boldsymbol{\beta}_{\texttt{lig}}, \boldsymbol{\beta}_{\texttt{trim}})$ is computed according to (17). With this, we then define the expected log-likelihood of new parameter estimates $\boldsymbol{\beta}'_{\texttt{lig}}$ and $\boldsymbol{\beta}'_{\texttt{trim}}$ given the current estimates $\boldsymbol{\beta}_{\texttt{lig}}$ and $\boldsymbol{\beta}_{\texttt{trim}}$ for a single sampled sequence $X$ as:

$$\begin{aligned}
&\ell(\boldsymbol{\beta}'_{\texttt{lig}}, \boldsymbol{\beta}'_{\texttt{trim}} \mid \boldsymbol{\beta}_{\texttt{lig}}, \boldsymbol{\beta}_{\texttt{trim}}; X, Q, I=0) \\
&\quad = \sum_{(t,m) \in A_X} P_{\text{annot}}(T=t, M=m \mid S=s, Q, I=0; X, \boldsymbol{\beta}_{\texttt{lig}}, \boldsymbol{\beta}_{\texttt{trim}}) \\
&\qquad\qquad \times \log P(T=t, M=m \mid S=s, Q, I=0; \boldsymbol{\beta}'_{\texttt{lig}}, \boldsymbol{\beta}'_{\texttt{trim}})
\end{aligned} \tag{19}$$

where $P_{\text{annot}}(T=t, M=m \mid S=s, Q=q, I=0; X, \boldsymbol{\beta}_{\texttt{lig}}, \boldsymbol{\beta}_{\texttt{trim}})$ and $P(T=t, M=m \mid S=s, Q=q, I=0; \boldsymbol{\beta}'_{\texttt{lig}}, \boldsymbol{\beta}'_{\texttt{trim}})$ are defined in (18) and (17), respectively. Similarly, we define the log-likelihood function for a random sample of observed sequences $\mathcal{X}$ as:

$$\mathcal{L}(\boldsymbol{\beta}'_{\texttt{lig}}, \boldsymbol{\beta}'_{\texttt{trim}} \mid \boldsymbol{\beta}_{\texttt{lig}}, \boldsymbol{\beta}_{\texttt{trim}}; \mathcal{X}, Q, I=0) = \sum_{X \in \mathcal{X}} C(X) \times \ell(\boldsymbol{\beta}'_{\texttt{lig}}, \boldsymbol{\beta}'_{\texttt{trim}} \mid \boldsymbol{\beta}_{\texttt{lig}}, \boldsymbol{\beta}_{\texttt{trim}}; X, Q, I=0) \tag{20}$$

where $C(X)$ represents the observed count of a specific sequence $X \in \mathcal{X}$ in the sampled data and $\ell(\boldsymbol{\beta}'_{\texttt{lig}}, \boldsymbol{\beta}'_{\texttt{trim}} \mid \boldsymbol{\beta}_{\texttt{lig}}, \boldsymbol{\beta}_{\texttt{trim}}; X, Q, I=0)$ is defined as in (19). The calculation of this expectation $\mathcal{L}$ constitutes the E-step of our EM procedure. Subsequently, in the minimization step (M-step), we update the model parameters by minimizing the negative log-likelihood of the observed data obtained in the E-step. This minimization step is performed using gradient descent with the `JAX` and `JAXopt` packages in Python [2, 1]. The algorithm iterates between the E and M steps until changes in the negative log-likelihood between successive iterations fall below a predefined threshold, indicating convergence.

#### Evaluating model using simulated data

To ensure our model returned expected outputs, we designed a data simulator capable of generating data under specific microhomology regimes. The simulator first samples a V-gene and J-gene according to IGoR-derived gene usage probabilities. Next, we establish probabilities for each trimming configuration using outputs from a version of our model that excludes microhomology terms, incorporating only trimming motif and base count terms. We then adjust these trimming probabilities using a tunable parameter to simulate the effect of microhomology, and the simulator samples a trimming configuration based on these adjusted probabilities. Finally, the simulator samples a ligation configuration uniformly, unless adjusted for microhomology effects by another tunable parameter. These two tunable parameters allow us to control the influence of microhomology on trimming and ligation choices. This process generates an observed simulated sequence, and by repeating it, we obtain a large set of simulated sequences to train and evaluate our model. We ran the simulator in four modes:

1. **No microhomology effect:** Both tunable microhomology parameters set to zero; microhomology does not influence trimming or ligation choices.
2. **Microhomology affects both trimming and ligation:** Both tunable microhomology parameters set to nonzero, positive values; microhomology increases probabilities for both trimming and ligation choices.
3. **Microhomology affects trimming, but not ligation:** Trimming-related parameter set to a nonzero, positive value and ligation-related parameter set to zero; microhomology increases trimming choice probabilities but does not affect ligation probabilities.
4. **Microhomology affects ligation, but not trimming:** Ligation-related parameter set to a nonzero, positive value and trimming-related parameter set to zero; microhomology increases ligation choice probabilities but does not affect trimming probabilities.

Using these simulated datasets, we trained our model to ensure that the expected inferred parameters were obtained. We also adjusted the strength of the microhomology-related effects using these tunable parameters to ensure our model could capture these signals.

#### Exploring the relationship between microhomology and trimming probabilities, independent of ligation

To quantify the effect of microhomology on trimming configuration probabilities independently of ligation, we restrict our training dataset to non-productive sequences *containing* N-insertions, as their presence suggests that germline-microhomology-mediated ligation did not occur. We aim to determine the influence of various sequence-level parameters on $P(R \mid S, Q, I > 0)$, where $R$ represents the IGoR-inferred trimming configuration, $S$ represents the gene pair, $Q$ represents the sequence productivity, and $I > 0$ represents nonzero N-insertions.

We previously demonstrated that local nucleotide identity surrounding trimming sites (the "trimming motif") and the counts of GC or AT nucleotides beyond these motifs (the "two-side base-count beyond") are highly predictive of trimming probabilities for single gene sequences [9]. Here, we model the probabilities of trimming configurations for gene pairs using these established parameters along with a new parameter related to "intermediate microhomology." This new parameter measures the importance of the average number of microhomologous nucleotides between two trimmed sequences, which, notably, are not homologous in the final rearranged sequence, see following section for definition. A summary of these model parameters and their corresponding weights for an arbitrary gene pair $S = (S_V, S_J)$ and trimming configuration $R = (R_V, R_J)$ is given in Table S2.

Using these model features, we define a weight function $f$ such that our model of $P(R \mid S, Q, I > 0)$ will be a normalized version of this weight. We parameterize $f$ using $\boldsymbol{\beta}$, the set of all model parameters, as follows:

$$\begin{aligned} f(R, S; \boldsymbol{\beta}) &:= f(R, S; \boldsymbol{\beta}_V^{\texttt{motif}}, \boldsymbol{\beta}_J^{\texttt{motif}}, \boldsymbol{\beta}_V^{\texttt{AT}}, \boldsymbol{\beta}_V^{\texttt{GC}}, \boldsymbol{\beta}_J^{\texttt{AT}}, \boldsymbol{\beta}_J^{\texttt{GC}}, \beta^{\texttt{iMH}}) \\ &:= f_{\texttt{motif}}(R_V, S_V; \boldsymbol{\beta}_V^{\texttt{motif}}) + f_{\texttt{motif}}(R_J, S_J; \boldsymbol{\beta}_J^{\texttt{motif}}) \\ &\quad + f_{\texttt{count}}(R_V, S_V; \boldsymbol{\beta}_V^{\texttt{AT}}, \boldsymbol{\beta}_V^{\texttt{GC}}) + f_{\texttt{count}}(R_J, S_J; \boldsymbol{\beta}_J^{\texttt{AT}}, \boldsymbol{\beta}_J^{\texttt{GC}}) \\ &\quad + f_{\texttt{iMH}}(R_V, R_J, S_V, S_J; \beta^{\texttt{iMH}}). \end{aligned} \tag{21}$$

Here, the parameter-specific weights $f_{\texttt{motif}}$, $f_{\texttt{count}}$, and $f_{\texttt{iMH}}$ are summarized in Table S2 and in the following section.

With this weight formulation, we can fit a conditional logit model which posits

$$P(R \mid S, Q, I > 0; \boldsymbol{\beta}) := \frac{\exp\big(f(R, S; \boldsymbol{\beta})\big)}{\sum_{R' \in \mathcal{R}} \exp\big(f(R', S; \boldsymbol{\beta})\big)}. \tag{22}$$

Here, $S$ and $R$ are random variables representing the V-gene and J-gene pair and trimming configuration, respectively, and $\mathcal{R}$ is the set of all reasonable trimming configurations.

The likelihood function $\ell(\boldsymbol{\beta})$ for a random sample of sequences is the likelihood of the model parameters $\boldsymbol{\beta}$ given a set of observed trimming configurations for specific gene pairs. The log likelihood function is:

$$\log \ell(\boldsymbol{\beta}) \\ = \sum_{S' \in \mathcal{S}} \sum_{R' \in \mathcal{R}} C(R', S', Q, I > 0) \cdot \log P(R' \mid S', Q, I > 0; \boldsymbol{\beta}) \tag{23}$$

where $\mathcal{S}$ represents the set of all gene pairs and $\mathcal{R}$ represents the set of all reasonable trimming configurations. Here, $C(R', S', Q, I > 0)$ is the count of sequences with trimming configuration $R'$, gene pair $S'$, nonzero N-insertions, and sequence productivity $Q$ and $P(R' \mid S', Q, I > 0; \boldsymbol{\beta})$ is given by Equation (22).

We include an additional regularization term for the intermediate-microhomology-specific parameters to help prevent over-fitting during model training. As such, we define a log loss function as

$$\mathcal{L}(\boldsymbol{\beta}) = -\log \ell(\boldsymbol{\beta}) + \lambda \cdot (\beta^{\texttt{iMH}})^2 \tag{24}$$

where $\log \ell(\boldsymbol{\beta})$ is given by (23) and $\lambda$ is a L2 regularization hyperparameter. We minimize this log loss function using gradient descent with the `JAX` and `JAXopt` packages in Python [2, 1]. We use a grid search to optimize the L2 regularization hyperparameter, $\lambda$. Notably, training this model without regularization (i.e. $\lambda = 0$) yields the same results as using the `mclogit` package in R, another implementation of conditional logistic regression that does not allow for regularization.

#### Defining "intermediate microhomology" parameters

In addition to the previously defined parameters, we define new microhomology-related parameters to model possible intermediate microhomology-mediated effects on the observed trimming configuration. We use the term "intermediate" because these nucleotides, while not directly participating in the final ligation, may temporarily influence intermediate steps such as trimming. Let $a$ be a non-negative integer value that represents the number of nucleotides 5' of each trimming site which are allowed to overlap between the two sequences when orienting the top strand of the V-gene sequence 5'-to-3' and the bottom strand of the J-gene sequence 3'-to-5' (e.g. as highlighted in yellow in Figure S10). Given a value of $a$, random variables $S_V$ and $S_J$ representing a V-gene and J-gene which each define a V-gene and J-gene sequence (both oriented 3'-to-5' as ordered lists), and random variables $R_V$ and $R_J$ representing V-gene and J-gene trimming amounts, the sub-sequences corresponding to this overlapping region are defined by the following ordered lists

$$\mathsf{seq}_{\texttt{Vmh}}(S_V, R_V, a) = \begin{cases} \big(S_V(R_V + 2 - j)\big)_{j=(1-a)}^{0} & \text{if } a \geq 1 \\ () & \text{if } a = 0 \end{cases} \tag{25}$$

and

$$\mathsf{seq}_{\texttt{Jmh}}(S_J, R_J, a) = \begin{cases} \big(S_J(R_J + 2 - j)\big)_{j=0}^{(1-a)} & \text{if } a \geq 1 \\ () & \text{if } a = 0. \end{cases} \tag{26}$$

Here, $S_V(R_V + 2 - j)$ and $S_J(R_J + 2 - j)$ represent the nucleotide identities at a sequence position $j$ where positions $j \leq 0$ represent sequence positions 5' of the trimming sites and positions $j > 0$ represent sequence positions 3' of the trimming sites. The resulting sub-sequences, $\mathsf{seq}_{\texttt{Vmh}}(S_V, R_V, a)$ and $\mathsf{seq}_{\texttt{Jmh}}(S_J, R_J, a)$, are oriented in the 5'-to-3' and 3'-to-5' directions, respectively, making them complementary. To quantify microhomology, we can define a function $g$ which will count the number of complementary (i.e. microhomologous) nucleotides between two arbitrary overlapping, equal-length sequence regions, $x$ and $y$, as

$$g(x, y) = \sum_{i=0}^{\text{len}(x)} \begin{cases} 1 & \text{if } x(i) \text{ is complementary to } y(i) \\ 0 & \text{otherwise.} \end{cases} \tag{27}$$

It has been established that classical non-homologous end joining, which is the joining process used during V(D)J recombination, may involve up to four nucleotides of microhomology [5]. As such, we quantify the average number of non-contiguous microhomologous nucleotides across these overlapping interior subsequences corresponding to each $a \in \{1, 2, 3, 4\}$ as follows:

$$m(S_V, S_J, R_V, R_J) := \frac{\sum_{a \in \{1,2,3,4\}} g\big(\mathsf{seq}_{\mathtt{Vmh}}(S_V, R_V, a), \mathsf{seq}_{\mathtt{Jmh}}(S_J, R_J, a)\big)}{4} \tag{28}$$

With this average number of microhomologous nucleotides, we define an *intermediate microhomology* model parameter, $\beta^{\mathtt{iMH}}$, and a corresponding weight function for a pair of trimming sites $R_V$ and $R_J$ and genes $S_V$ and $S_J$:

$$f_{\mathtt{iMH}}(R_V, R_J, S_V, S_J; \beta^{\mathtt{iMH}}) := \beta^{\mathtt{iMH}} \cdot m(S_V, S_J, R_V, R_J) \tag{29}$$

using the previously defined sequences and the function $m$ as defined in (28).

#### 2 Supporting Figures and Tables

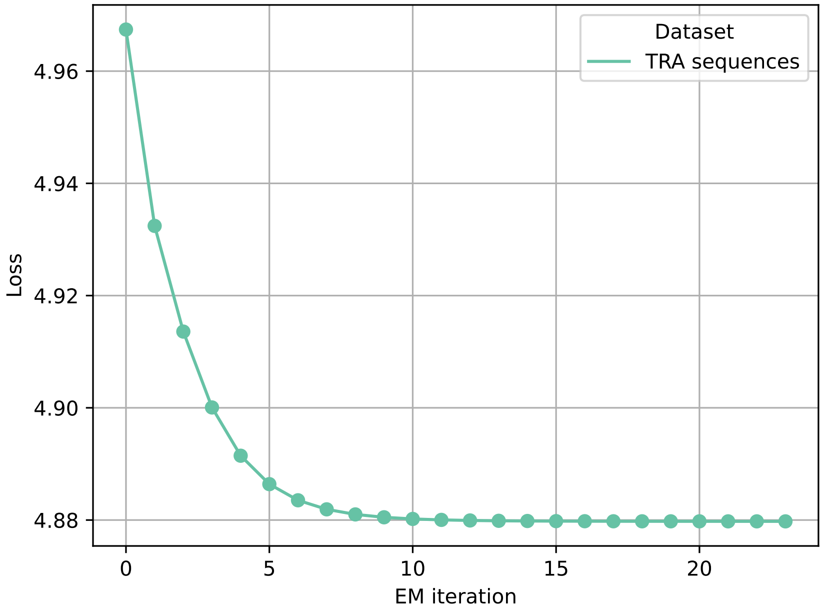


Figure S1: Convergence of the expectation-maximization (EM) algorithm using a training dataset of non-productive TCR$\alpha$ sequences without N-insertions, alongside their corresponding sets of potential microhomology-adapted annotations (as detailed in Methods). The y-axis represents the expected per-sequence log loss, as defined in (20).

Table S1: Summary of all notation used in our modeling.

| **Variable** | **Description** |
|---|---|
| **General notation** | |
| $X$ | arbitrary sampled sequence |
| $\mathcal{X}$ | set of sampled sequences |
| $S_V, S_J$ | random variables for the V- and J-gene |
| $S$ | ordered pair of genes, $(S_V, S_J)$ |
| $R_V, R_J$ | random variables for nucleotides deleted from the V/J-gene, inferred by IGoR |
| $R$ | ordered pair of IGoR-inferred trimming amounts, $(R_V, R_J)$ |
| $I$ | random variable for number of N-insertions |
| $Q$ | deterministic variable for sequence productivity |
| $M$ | random variable for number of microhomologous nucleotides within the observed sequence; also referred to as a "ligation configuration" |

| | |
|---|---|
| $T_V, T_J$ | random variables for microhomology-adapted trimming amounts from V/J-gene |
| $T$ | pair of microhomology-adapted trimming amounts, $(T_V, T_J)$; also referred to as a "trimming configuration" |
| $A_X$ | set of possible microhomology-adapted trimming and ligation annotations for a sequence $X$ |

**Motif parameter notation**

| | |
|---|---|
| $\boldsymbol{\beta}_V^{\texttt{motif}}, \boldsymbol{\beta}_J^{\texttt{motif}}$ | set of all V/J-gene motif parameters, defined in detail within Supplementary Materials |
| $f_{\texttt{motif}}(T_V, S_V; \boldsymbol{\beta}_V^{\texttt{motif}})$ | V-gene motif weight function (5) |
| $f_{\texttt{motif}}(T_J, S_J; \boldsymbol{\beta}_J^{\texttt{motif}})$ | J-gene motif weight function (5) |

**Base-count-beyond parameter notation**

| | |
|---|---|
| $\boldsymbol{\beta}_V^{\texttt{AT}}, \boldsymbol{\beta}_V^{\texttt{GC}}$ | V-gene AT/GC base-count-beyond parameters, defined in detail within Supplementary Materials |
| $\boldsymbol{\beta}_J^{\texttt{AT}}, \boldsymbol{\beta}_J^{\texttt{GC}}$ | J-gene AT/GC base-count-beyond parameters |
| $f_{\texttt{count}}(T_V, S_V; \boldsymbol{\beta}_V^{\texttt{AT}}, \boldsymbol{\beta}_V^{\texttt{GC}})$ | V-gene base-count-beyond weight function (10) |
| $f_{\texttt{count}}(T_J, S_J; \boldsymbol{\beta}_J^{\texttt{AT}}, \boldsymbol{\beta}_J^{\texttt{GC}})$ | J-gene base-count-beyond weight function (10) |

**Microhomology parameter notation**

| | |
|---|---|
| $\beta^{\texttt{trimMH}}, \beta^{\texttt{ligMH}}$ | trimming/ligation microhomology parameters |
| $f_{\texttt{trimMH}}(T, S; \beta^{\texttt{trimMH}})$ | trimming-related microhomology weight function (11) |
| $f_{\texttt{ligMH}}(M; \beta^{\texttt{ligMH}})$ | ligation-related microhomology weight function (12) |

**Model notation**

| | |
|---|---|
| $\boldsymbol{\beta}_{\texttt{trim}}$ | set of all trimming-related model parameters: $\boldsymbol{\beta}_V^{\texttt{motif}}, \boldsymbol{\beta}_J^{\texttt{motif}}, \boldsymbol{\beta}_V^{\texttt{AT}}, \boldsymbol{\beta}_J^{\texttt{AT}}, \boldsymbol{\beta}_V^{\texttt{GC}}, \boldsymbol{\beta}_J^{\texttt{GC}}, \beta^{\texttt{trimMH}}$ |
| $\boldsymbol{\beta}_{\texttt{lig}}$ | set of all ligation-related model parameters: $\beta^{\texttt{ligMH}}$ |
| $f_{\texttt{trim}}(T, S; \boldsymbol{\beta}_{\texttt{trim}})$ | trimming-related weight function (2) |
| $f_{\texttt{lig}}(T, M, S; \boldsymbol{\beta}_{\texttt{lig}})$ | ligation-related weight function (3) |
| $P(T, M \mid S, Q, I = 0; \boldsymbol{\beta}_{\texttt{trim}}, \boldsymbol{\beta}_{\texttt{lig}})$ | two-step conditional logit model (17) |
| $P_{\text{annot}}(T, M \mid S, Q, I = 0; X, \boldsymbol{\beta}_{\texttt{lig}}, \boldsymbol{\beta}_{\texttt{trim}})$ | model-derived trimming and ligation annotation probability (18) |
| $\ell(\boldsymbol{\beta}'_{\texttt{lig}}, \boldsymbol{\beta}'_{\texttt{trim}} \mid \boldsymbol{\beta}_{\texttt{lig}}, \boldsymbol{\beta}_{\texttt{trim}}; X, Q, I = 0)$ | expected log-likelihood for single sequence (19) |
| $\mathcal{L}(\boldsymbol{\beta}'_{\texttt{lig}}, \boldsymbol{\beta}'_{\texttt{trim}} \mid \boldsymbol{\beta}_{\texttt{lig}}, \boldsymbol{\beta}_{\texttt{trim}}; \mathcal{X}, Q, I = 0)$ | log-likelihood for observed sequences (20) |

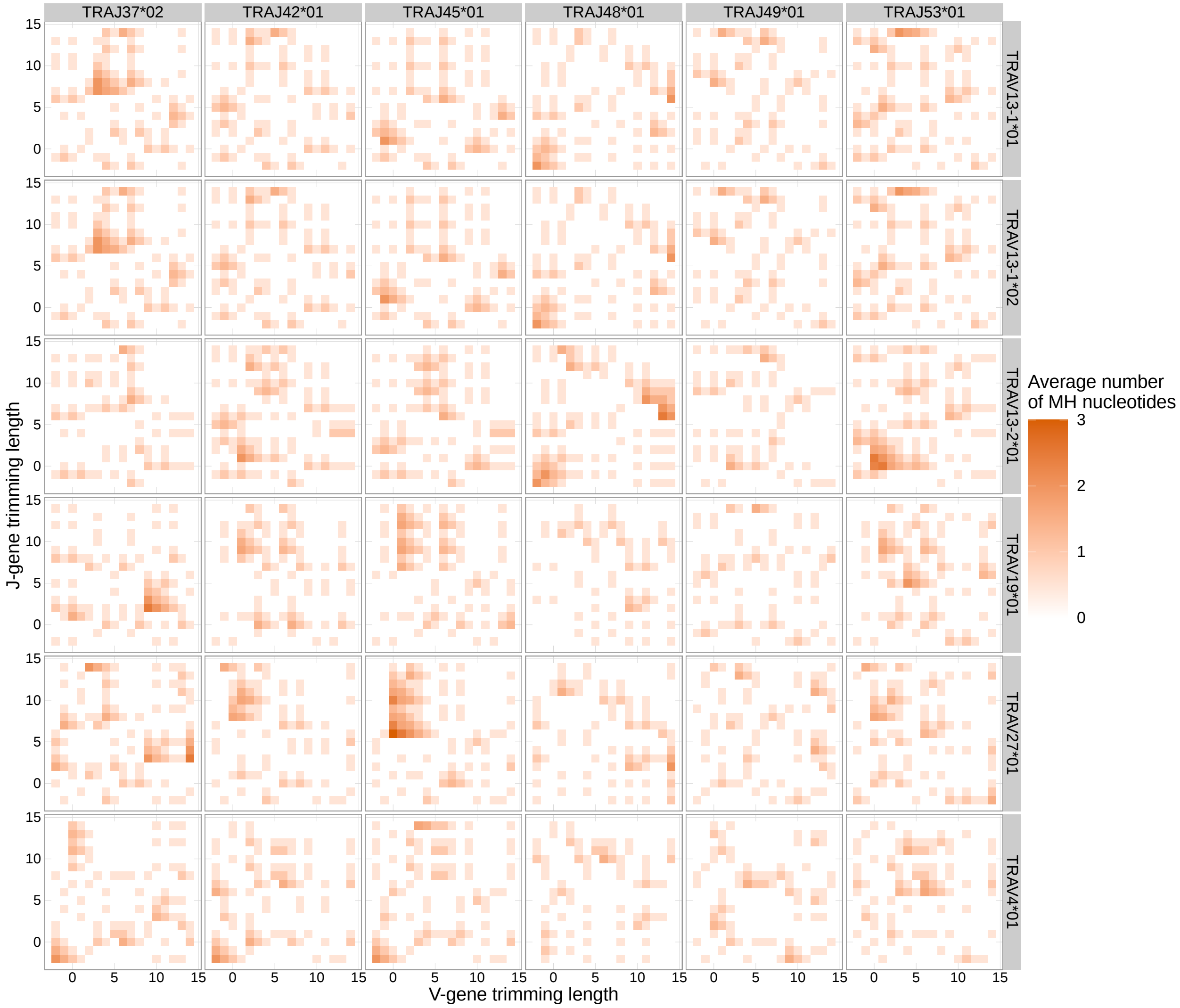


Figure S2: The distribution of complementary sequence regions capable of forming microhomologous regions during V(D)J recombination varies by trimming amounts and V-J gene pairs. Depending on the gene pair, there is potential for both interior and terminal microhomology (MH). For instance, the TRAV13-1*01 and TRAJ48*01 gene pair shows potential for terminal microhomology (e.g. the average number of MH nucleotides for the untrimmed sequences—both genes trimmed at the -2 site—is nonzero), as well as interior microhomology. In contrast, the TRAV13-1*01 and TRAJ37*02 gene pair lacks terminal microhomology potential (e.g. the average number of MH nucleotides for the untrimmed sequences is zero) but has an abundance of interior microhomology. The average microhomology counts are calculated across all possible ligation configurations for each gene pair trimming configuration. Only the most frequently used gene pairs are plotted here.

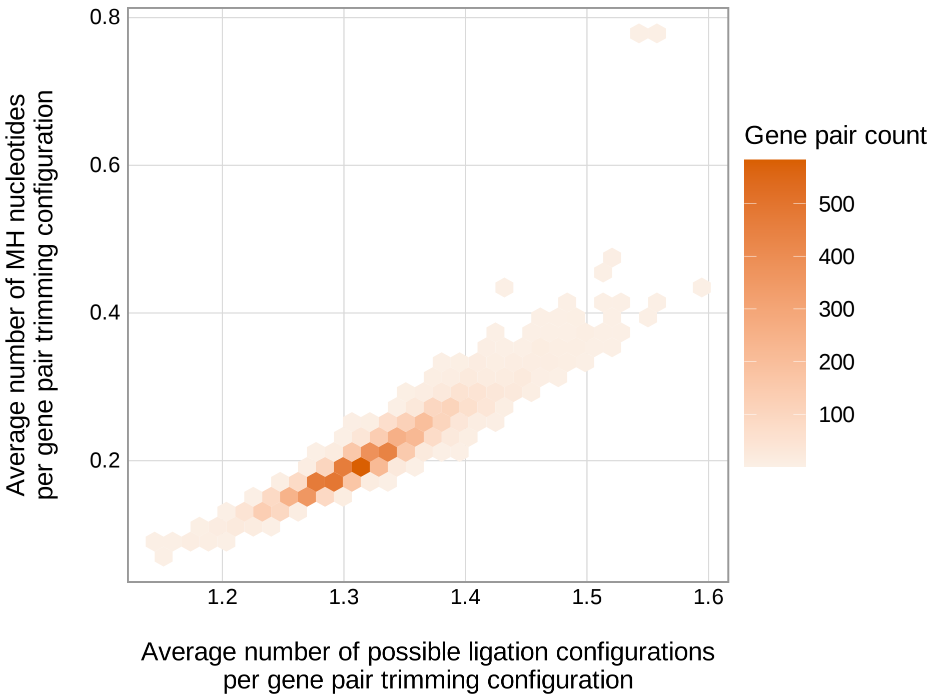


Figure S3: Complementary sequence regions capable of forming microhomologous regions during V(D)J recombination are common between germline V- and J-genes in the *TRA* locus. As the average number of microhomologous nucleotides increases for a given gene pair trimming configuration, so does the average number of possible ligation configurations. The median average number of microhomologous nucleotides across all possible gene pair trimming configurations is 0.1978, corresponding to a median of 1.3149 possible ligation configurations. Since a median of exactly one ligation configuration would indicate that all configurations involve zero microhomology, this suggests that most trimming configurations result in multiple ligation outcomes—both with and without microhomology. The average values are calculated across all trimming configurations for each gene pair, with each gene pair plotted only once.

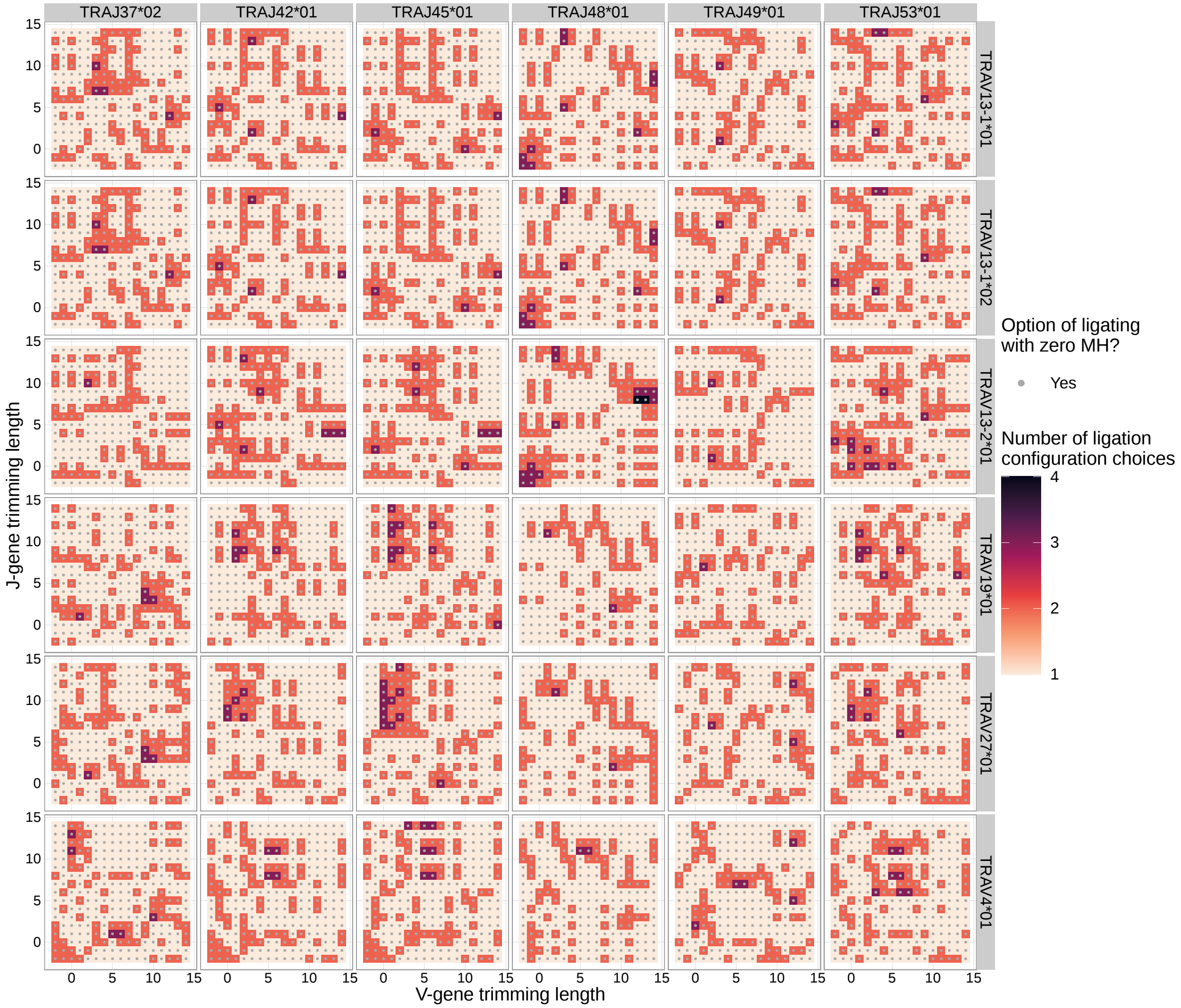


Figure S4: The distribution of trimming configurations with multiple ligation options (e.g. varying amounts of microhomology in the observed sequence) varies across V-J gene pairs. All gene pair trimming configurations can ligate with zero nucleotides of microhomology (indicated by gray dots in the plot), providing at least one ligation configuration choice. Depending on the gene pair, there may be potential for both interior- and terminal-microhomology-mediated trimming and ligation (e.g. ligation using nonzero microhomology), which would increase the number of possible ligation configuration choices. For example, the TRAV13-1*01 and TRAJ48*01 gene pair shows potential for terminal-microhomology-mediated ligation (e.g. ligating the untrimmed sequences—both genes trimmed at the -2 site—using nonzero microhomology), as well as interior-microhomology-mediated ligation. In contrast, the TRAV13-1*01 and TRAJ37*02 gene pair lacks potential for terminal-microhomology-mediated ligation (e.g. the untrimmed sequences can only be ligated using zero microhomology) but has substantial potential for interior-microhomology-mediated ligation. Only the most frequently used gene pairs are plotted here.

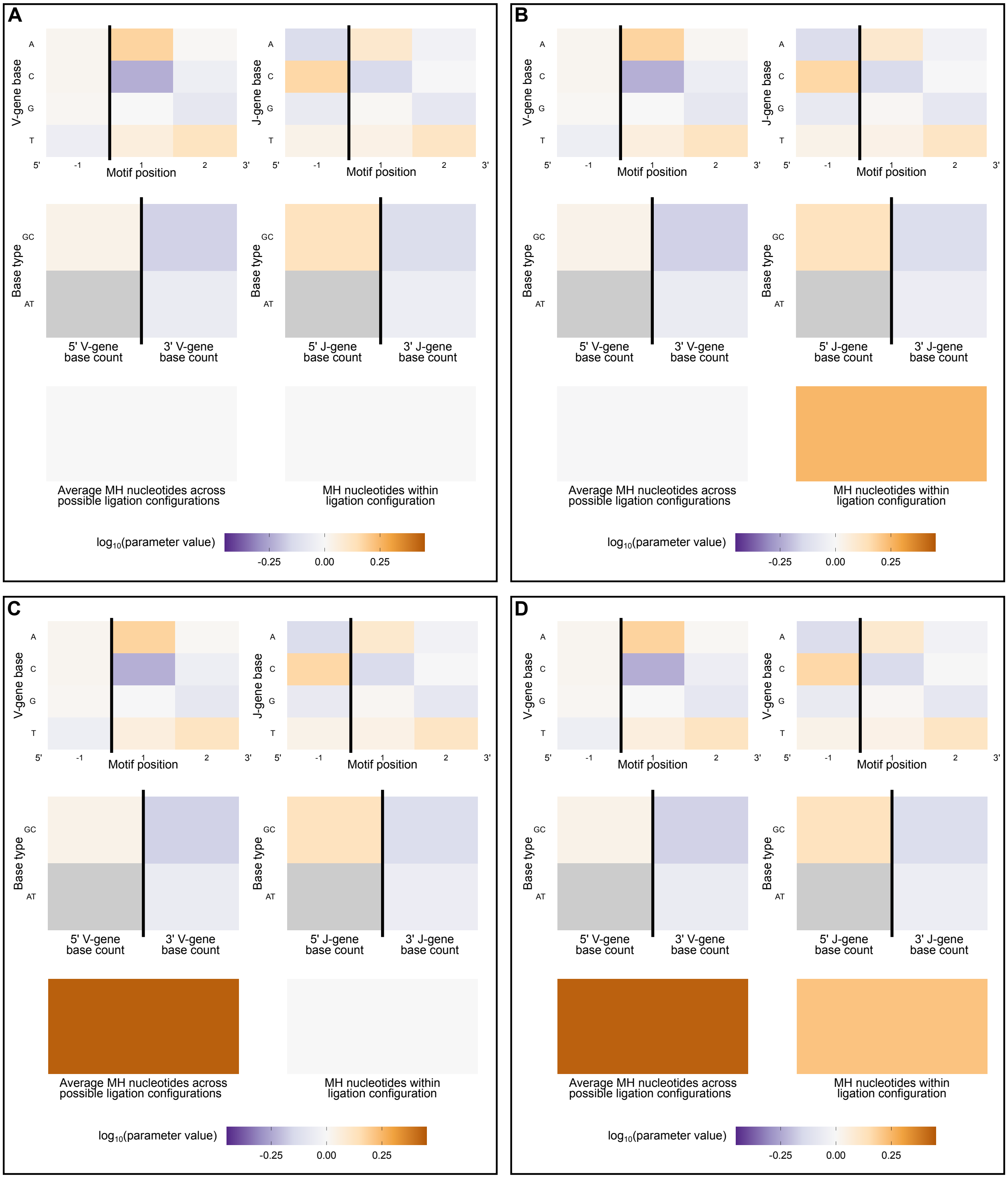


Figure S5: Parameters inferred from simulated data emulating varying levels of microhomology involvement in V(D)J recombination trimming and ligation (see Supplementary Materials). The model was trained using four different simulated datasets. (**A**) Parameters inferred from data where microhomology does not influence trimming or ligation choices. In this scenario, the microhomology-related parameters show no signal. (**B**) Parameters inferred from data where microhomology increases ligation probabilities but does not affect trimming probabilities. Here, the trimming-related microhomology parameter shows no signal, while the ligation-related parameter shows a strong positive signal. (**C**) Parameters inferred from data where microhomology increases trimming probabilities but does not affect ligation probabilities. In this case, the ligation-related microhomology parameter shows no signal, while the trimming-related parameter shows a strong positive signal. (**D**) Parameters inferred from data where microhomology increases both trimming and ligation probabilities. As expected, both microhomology-related parameters exhibit strong positive signals. The patterns observed in this simulated dataset closely matches those inferred from actual data. All trimming motif and two-side base count parameters remain consistent across all simulated datasets.

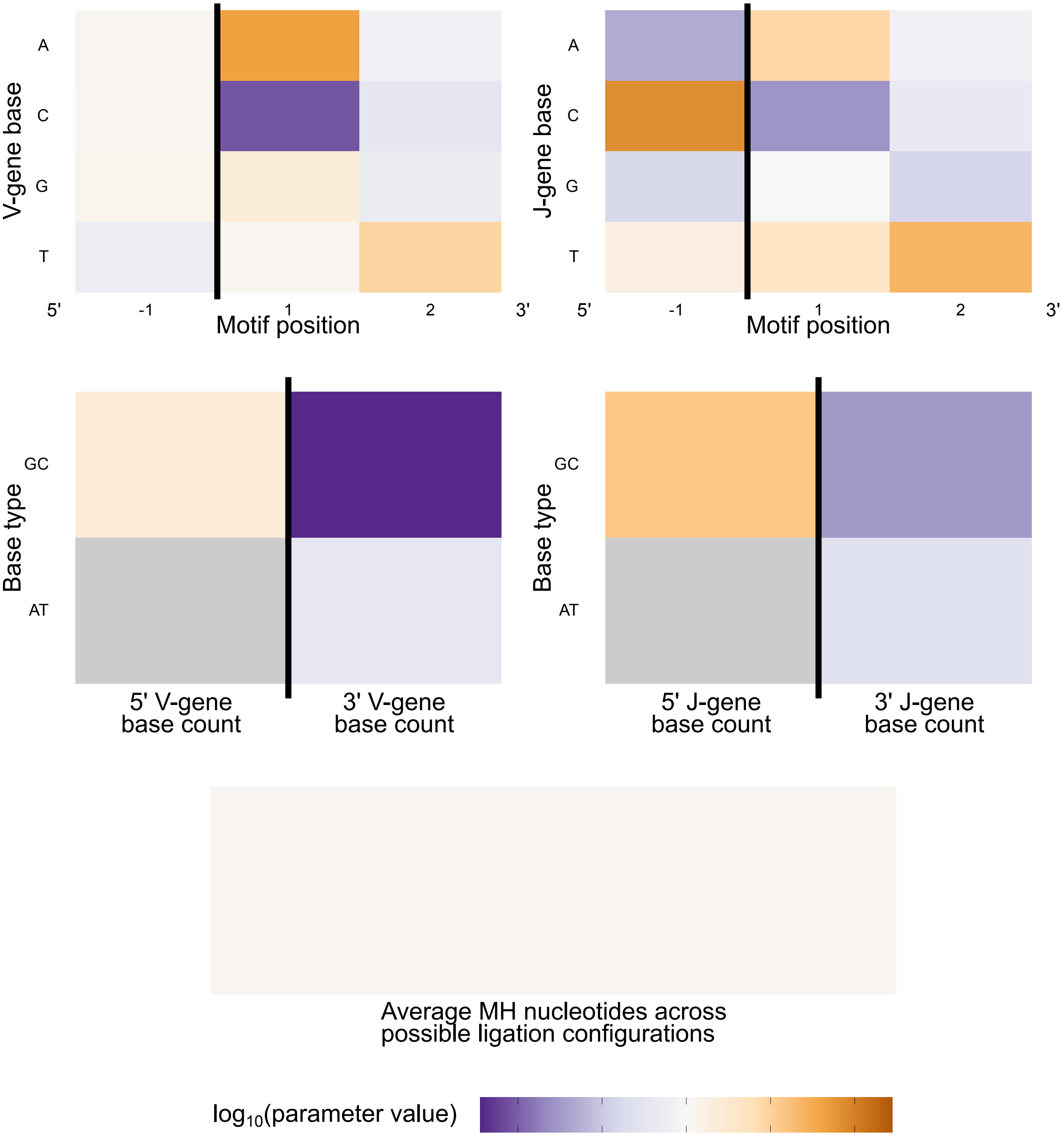


Figure S6: Parameters inferred from a model trained on sequences with N-insertions, which lack germline-dependent ligation. Due to the unknown composition of inserted nucleotides prior to ligation, ligation patterns cannot be detected, allowing this model to specifically explore trimming probabilities independent of ligation (see Supplementary Materials). The ligation-related microhomology parameter was excluded from this model. Inferred trimming motif and two-side base count parameters align with previous analyses of individual V- and J-gene sequences [9]. The inferred trimming-related microhomology parameter indicates that microhomology has only a minimal effect on trimming probabilities, suggesting a limited independent role of microhomology in trimming.

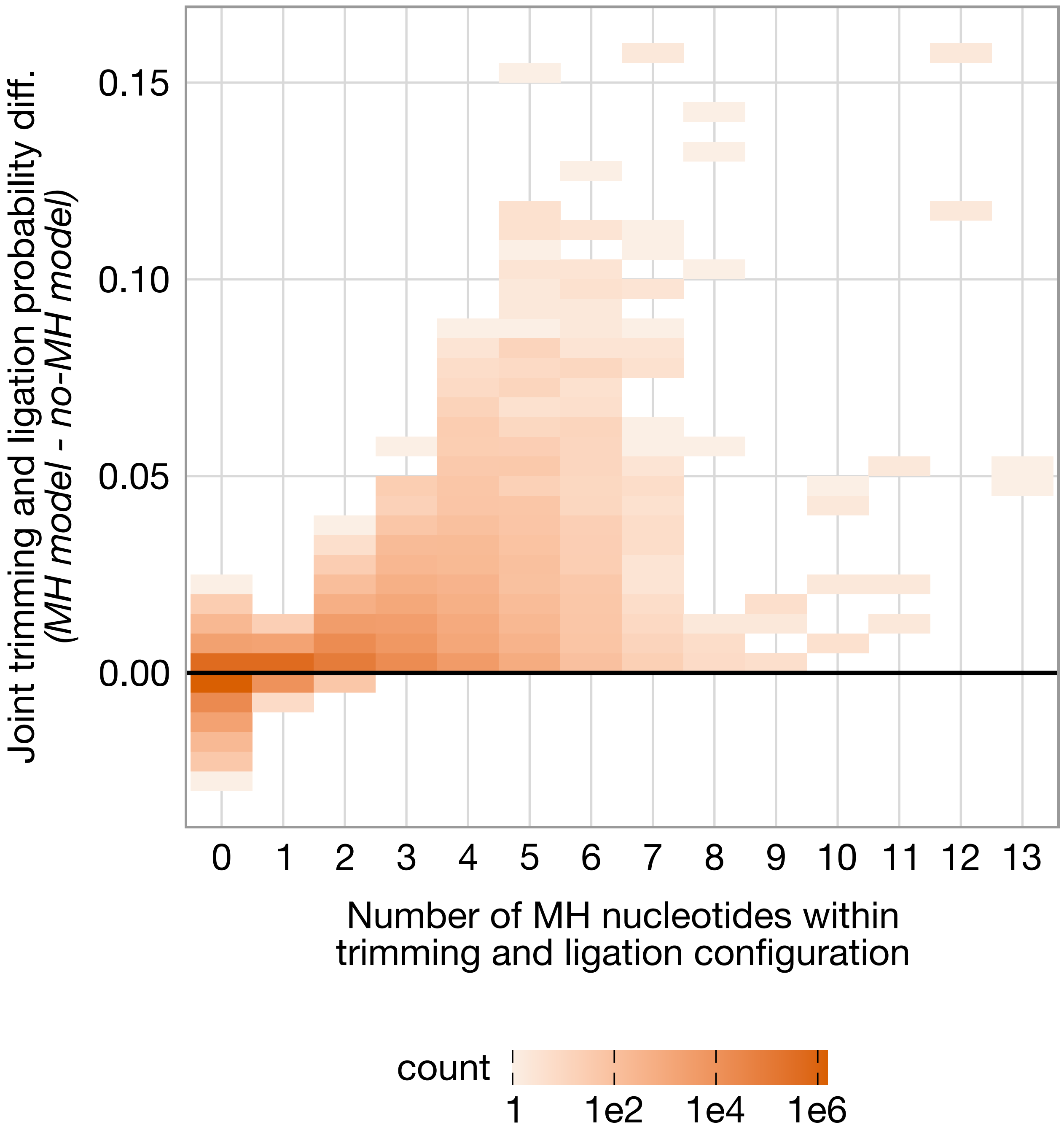


Figure S7: As the number of microhomologous nucleotides in a trimming and ligation configuration increases, the difference in joint trimming and ligation probabilities between the model parameterizing microhomology (MH model) and the one that does not (no-MH model) becomes larger. Both models include trimming motif and base-count parameters and are identical except for the inclusion of microhomology terms. The plotted differences are calculated using probabilities normalized across all possible trimming and ligation configurations for each V-J gene pair, ensuring the sum of these differences is zero.

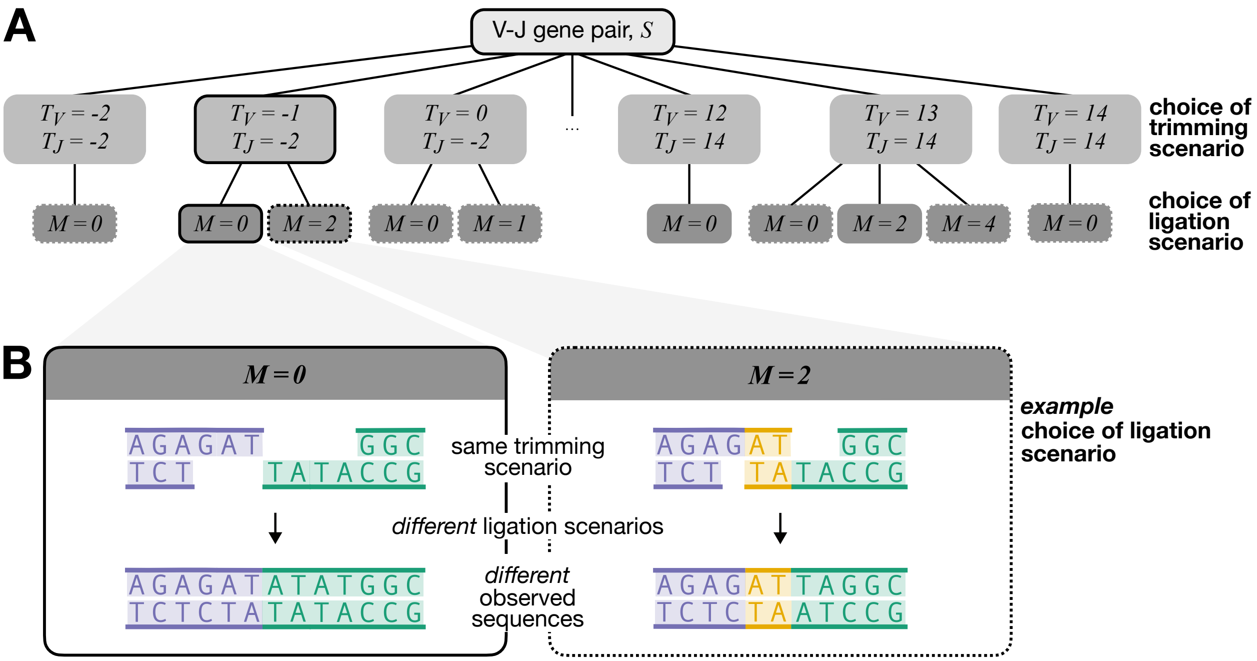


Figure S8: Version of Figure 5 specifying variable notation. **(A)** Schematic of trimming and ligation choices for an arbitrary V-J gene pair, denoted as an ordered pair $S = (S_V, S_J)$, where $S_V$ and $S_J$ are random variables representing V- and J-genes, respectively. The first choice is the trimming configuration, represented as the ordered pair $T = (T_V, T_J)$ where $T_V$ and $T_J$ are random variables denoting the microhomology-adapted trimming amounts for the V- and J-genes (e.g. each gene can be trimmed by -2 to 14 nucleotides). The next choice is the ligation configuration, represented by the random variable $M$, which captures the number of microhomologous nucleotides used. The available ligation configuration choices will depend on the germline sequences of the two genes being joined. Trimming and ligation configurations resulting in productive and nonproductive sequences are shown in solid and dashed boxes, respectively. **(B)** Illustration of the possible ligation configurations for an example pair of trimmed sequences. The chosen ligation configuration affects the resulting observed sequence. The trimmed V-gene sequence is shown in purple, the trimmed J-gene sequence is shown in green, and microhomologous nucleotides are shown in yellow. Deletions are indexed such that a deletion of 0 corresponds to the end of the germline gene sequence (two P-nucleotides trimmed) and -2 corresponds to the end of the full sequence (no P-nucleotides or gene sequence nucleotides trimmed).

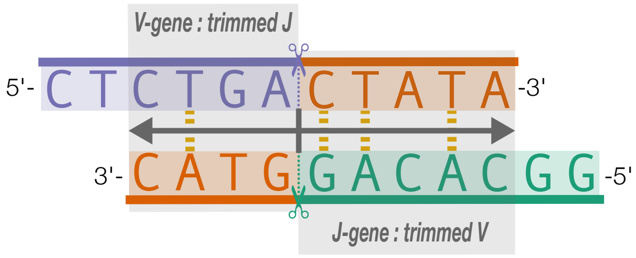


Figure S9: Cartoon showing the alignment of V-gene (purple) and J-gene (green) sequences at their inferred trimming sites (marked with scissors) without gaps. Trimmed regions of each sequence are shown in orange. We focus on two regions: (1) overlap between V-gene and trimmed J-gene (*V-gene:trimmed-J*) and (2) overlap between J-gene and trimmed V-gene (*J-gene:trimmed-V*). The function $h$ in (1) counts contiguous, complementary nucleotides from the aligned trimming site (indexed as zero). Arrows show the counting direction for each region. Contiguous, complementary nucleotides are counted only if adjacent to the trimming site. In this example, *V-gene:trimmed-J* has 0 contiguous complementary nucleotides, and *J-gene:trimmed-V* has 2.

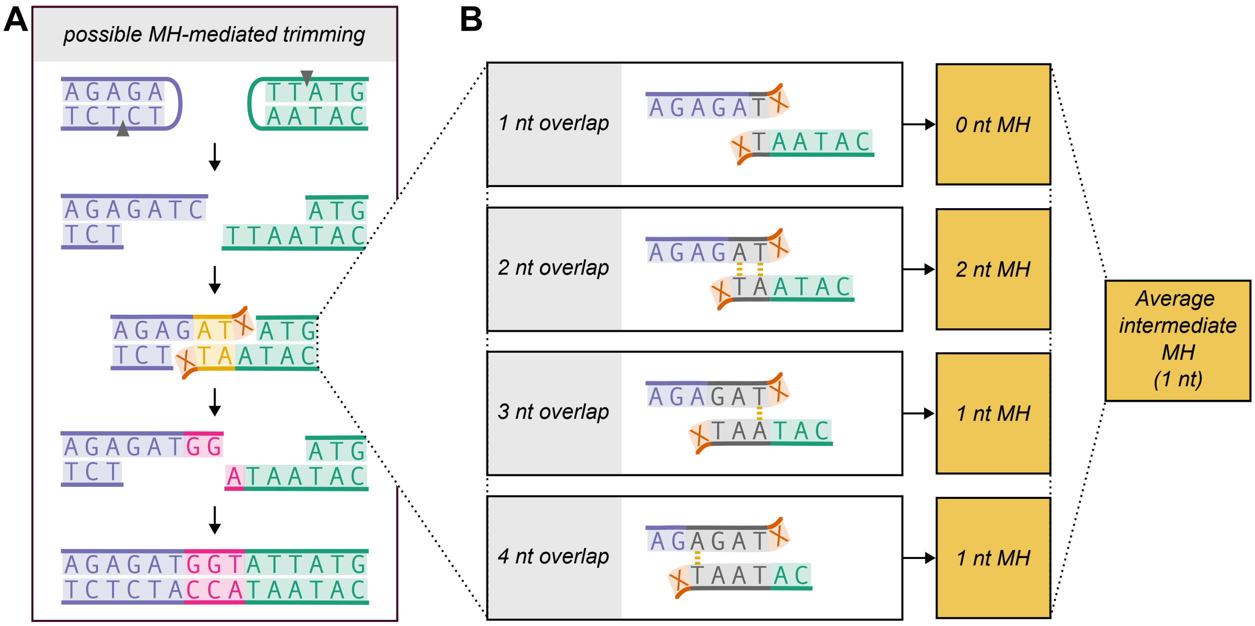


Figure S10: (**A**) Diagram illustrating V(D)J recombination steps, emphasizing possible internal/intermediate microhomology during trimming. This intermediate microhomology may occur in both single-stranded overhangs and double-stranded regions due to sequence breathing. The final joined sequence, post-N-insertion (pink), is shown in the last box. (**B**) Definition of overlapping sequence regions for specified V-gene, J-gene, and trimming configuration. The top strand of a V-gene and the bottom strand of a J-gene are each shown with one nucleotide removed (trimmed nucleotides in orange). Overlapping regions (highlighted in gray) are obtained by aligning the sequences such that an integer value, $a$, of nucleotides 5' of each trimming site overlap. The value of $a$ ranges from 1 to 4 nucleotides. Despite what is shown in this example, two genes can be trimmed by different amounts and still yield these overlapping regions. Complementary nucleotides in these regions are indicated by vertical yellow lines. The final joined sequence post-trimming and insertion is shown in the last box of panel (A).

Table S2: Summary of all parameters and parameter-specific weights for an arbitrary gene pair $S = (S_V, S_J)$ and trimming configuration $R = (R_V, R_J)$. Detailed definitions of each parameter and corresponding weights are located within the Supplementary Materials.

| **Parameter** | **Description** | **Notation** | **Parameter weight** |
|---|---|---|---|
| *V-gene motif* parameters | parameterizes the importance of several nucleotides on either side of the V-gene trimming site | $\boldsymbol{\beta}_V^{\texttt{motif}}$ | $f_{\texttt{motif}}(R_V, S_V; \boldsymbol{\beta}_V^{\texttt{motif}})$ (5) |
| *J-gene motif* parameters | parameterizes the importance of several nucleotides on either side of the J-gene trimming site | $\boldsymbol{\beta}_J^{\texttt{motif}}$ | $f_{\texttt{motif}}(R_J, S_J; \boldsymbol{\beta}_J^{\texttt{motif}})$ (5) |
| *V-gene base-count-beyond* parameters | parameterizes the importance of the counts of GC and AT nucleotides beyond the V-gene trimming motif | $\boldsymbol{\beta}_V^{\texttt{AT}}$ and $\boldsymbol{\beta}_V^{\texttt{GC}}$ | $f_{\texttt{count}}(R_V, S_V; \boldsymbol{\beta}_V^{\texttt{AT}}, \boldsymbol{\beta}_V^{\texttt{GC}})$ (10) |
| *J-gene base-count-beyond* parameters | parameterizes the importance of the counts of GC and AT nucleotides beyond the J-gene trimming motif | $\boldsymbol{\beta}_J^{\texttt{AT}}$ and $\boldsymbol{\beta}_J^{\texttt{GC}}$ | $f_{\texttt{count}}(R_J, S_J; \boldsymbol{\beta}_J^{\texttt{AT}}, \boldsymbol{\beta}_J^{\texttt{GC}})$ (10) |
| *inter-mediate microhomol-ogy* parameters | parameterizes the importance of the average number of non-contiguous intermediate microhomology between a gene pair given a trimming configuration | $\beta^{\texttt{iMH}}$ | $f_{\texttt{iMH}}(R_V, R_J, S_V, S_J; \beta^{\texttt{iMH}})$ (29) |
| *trimming-related observed mi-crohomology* parameters | parameterizes the importance of the average number of contiguous microhomology across all possible ligation configurations for a given trimming configuration and gene pair | $\beta^{\texttt{trimMH}}$ | $f_{\texttt{trimMH}}(T, S; \beta^{\texttt{trimMH}})$ (11) |
| *ligation-related observed mi-crohomology* parameters | parameterizes the importance of the number of contiguous microhomologous nucleotides for a given ligation configuration, trimming configuration, and gene pair | $\beta^{\texttt{ligMH}}$ | $f_{\texttt{ligMH}}(M; \beta^{\texttt{ligMH}})$ (12) |

### 3 SI References


[1] Mathieu Blondel, Quentin Berthet, Marco Cuturi, Roy Frostig, Stephan Hoyer, Felipe Llinares-López, Fabian Pedregosa, and Jean-Philippe Vert. Efficient and modular implicit differentiation. *arXiv [cs.LG]*, May 2021.

[2] James Bradbury, Roy Frostig, Peter Hawkins, Matthew James Johnson, Chris Leary, Dougal Maclaurin, George Necula, Adam Paszke, Jake VanderPlas, Skye Wanderman-Milne, and Qiao Zhang. JAX: composable transformations of Python+NumPy programs, 2018.

[9] Magdalena L Russell, Noah Simon, Philip Bradley, and Frederick A Matsen, 4th. Statistical inference reveals the role of length, GC content, and local sequence in V(D)J nucleotide trimming. *Elife*, 12, May 2023.

[10] Santosh K Srivastava and Harlan S Robins. Palindromic nucleotide analysis in human T cell receptor rearrangements. *PLoS One*, 7(12):e52250, December 2012.
